# Translating from mice to humans: using preclinical blood-based biomarkers for the prognosis and treatment of traumatic brain injury

**DOI:** 10.1101/2023.12.01.569152

**Authors:** Ilaria Lisi, Federico Moro, Edoardo Mazzone, Niklas Marklund, Francesca Pischiutta, Firas Kobeissy, Xiang Mao, Frances Corrigan, Adel Helmy, Fatima Nasrallah, Valentina Di Pietro, Laura B Ngwenya, Luis Portela, Bridgette Semple, Douglas H. Smith, Cheryl Wellington, David J Loane, Kevin Wang, Elisa R Zanier, the InTBIR Fundamental & Translational Working Group

**Affiliations:** Department of Acute Brain and Cardiovascular Injury, Istituto di Ricerche Farmacologiche Mario Negri IRCCS, Milan, Italy; Department of Clinical Sciences Lund, Neurosurgery, Lund University and Skåne University Hospital, Lund, Sweden; Department of Neurobiology, Center for Neurotrauma, Multiomics & Biomarkers, Morehouse School of Medicine, Atlanta, GA, USA; Department of Neurosurgery, Beijing TianTan Hospital, Capital Medical University, Beijing, China; School of Biomedicine, Faculty of Health and Medical Sciences, The University of Adelaide, Australia; Division of Neurosurgery, Department of Clinical Neurosciences, University of Cambridge, UK; Queensland Brain Institute, The University of Queensland, Queensland, Australia; Institute of Inflammation and Ageing, College of Medical and Dental Sciences, University of Birmingham, Birmingham, UK; Department of Neurosurgery, University of Cincinnati College of Medicine, Cincinnati, OH, USA; Department of Biochemistry, ICBS, Federal University of Rio Grande do Sul – UFRGS, Porto Alegre, RS, Brasil; Department of Neuroscience, Central Clinical School, Monash University, Melbourne, VIC, Australia; Center for Brain Injury and Repair and the Department of Neurosurgery, Perelman School of Medicine, University of Pennsylvania, Philadelphia, PA, USA; Department of Pathology, Djavad Mowafaghain Centre for Brain Health, International Collaboration on Repair Discoveries, School of Biomedical Engineering, University of British Columbia. Canada.; School of Biochemistry and Immunology, Trinity College Dublin, Dublin, Ireland

## Abstract

Rodent models are important research tools for studying the pathophysiology of traumatic brain injury (TBI) and developing potential new therapeutic interventions for this devastating neurological disorder. However, the failure rate for the translation of drugs from animal testing to human treatments for TBI is 100%, perhaps due, in part, to distinct timescales of pathophysiological processes in rodents versus humans that impedes translational advancements. Incorporating clinically relevant biomarkers in preclinical studies may provide an opportunity to calibrate preclinical models to human TBI biomechanics and pathophysiology. To support this important translational goal, we conducted a systematic literature review of preclinical TBI studies in rodents measuring blood levels of clinically used NfL, t-Tau, p-Tau, UCH-L1, or GFAP, published in PubMed/MEDLINE up to June 13th, 2023. We focused on blood biomarker temporal trajectories and their predictive and pharmacodynamic value and discuss our findings in the context of the latest clinical TBI biomarker data. Out of 369 original studies identified through the literature search, 71 met the inclusion criteria, with a median quality score on the CAMARADES checklist of 5 (interquartile range 4-7). NfL was measured in 17 preclinical studies, GFAP in 41, t-Tau in 17, p-Tau in 7, and UCH-L1 in 19 preclinical studies. Data in rodent models show that all blood biomarkers exhibited injury severity-dependent elevations, with GFAP and UCH-L1 peaking within hours after TBI, NfL peaking within days after TBI and remaining elevated up to 6 months post-injury, whereas t-Tau and p-Tau levels were gradually increased many weeks after TBI. Blood NfL levels emerges as a prognostic indicator of white matter loss after TBI, while both NfL and GFAP hold promise for pharmacodynamic studies of neuroprotective treatments. Therefore, blood-based preclinical biomarker trajectories could serve as important anchor points that may advance translational research in the TBI field. However, further investigation into biomarker levels in the subacute and chronic phases will be needed to more clearly define pathophysiological mechanisms and identify new therapeutic targets for TBI.

## Introduction

Traumatic brain injury (TBI) is a leading cause of death and disability worldwide that affects 60 million individuals annually (1). With a global economic burden of approximately 400 billion USD each year (1) and 82,000 TBI-related deaths recorded annually in Europe alone (2), the impact of TBI is unmistakably significant. Indeed, TBI is a multifaceted and dynamic neurological condition, with the primary biomechanical injury to the brain triggering a series of secondary pathophysiological events including – but not limited to – neuronal and astroglial damage, blood brain barrier (BBB) disruption, and persistent inflammation. A major challenge is the astonishing heterogeneity of clinical TBI across severities, injury biomechanics and demographic components, which can affect secondary intracerebral and systemic responses, and long-term recovery processes. This heterogeneity poses a significant challenge to the effective translation of neurotherapeutic interventions for TBI. While animal models have proven invaluable to mitigate this inherent heterogeneity, there has been 100% failure in translating promising preclinical drug treatment strategies to the clinic (3). Hence, efforts are being redirected towards integrating clinically relevant measures into preclinical research to facilitate this transition and overcome this block in translation. Clinically, blood protein biomarkers have emerged as critical tools to investigate the pathophysiological mechanisms underlying secondary injury post-TBI, classify injury severity (4,5), track injury progression (6,7), guide clinical decision making (8), anticipate clinical outcomes (7,9–11), and monitor the responsiveness to treatment (12,13). Blood biomarker incorporation in preclinical TBI models may serve as an important calibration tool, enabling the selection of the most suitable experimental model that will optimally relate preclinical mechanistic studies with clinical observations. This alignment of clinically relevant biomarkers will enhance the likelihood of effectively translating neuroprotective treatments from laboratory experiments to real-world medical practices.

Below, we provide a brief introduction to key blood-based biomarkers currently being developed as diagnostic and prognostic indicators of brain injury severity and outcome in humans.

### Neurofilament Light (NfL)

Neurofilaments are structural proteins located in the neuronal cytoplasm that confer essential stability to neurons, particularly in neurons that have large myelinated axons (14). Among the neurofilaments, NfL is the most abundant and soluble constituent, and circulating NfL levels remain low under physiological conditions (15). Following a TBI, the biomechanical forces involved lead to significant axonal injury due to white matter tracts being highly susceptible to mechanical stress (3,16). Secondary injury can also lead to progressive and exacerbated axonal damage (17). Following acute axonal injury and/or chronic neurodegeneration, NfL is released into the interstitial fluid, which communicates freely with the cerebrospinal fluid (CSF) and the blood stream when the BBB is damaged. NfL is an established blood biomarker of axonal injury in several neurodegenerative conditions, including TBI (7,18,19). Notably, clinical investigations centered around moderate-to-severe TBI have documented a distinct sub-acute peak in plasma NfL levels, which remains persistently elevated many years after TBI in humans (7,12,20).

### Glial Fibrillary Acidic Protein (GFAP)

GFAP is an intermediate filament III protein expressed by astrocytes that plays a role in the maintenance of both astrocytic skeletal structures and the integrity of the BBB (21). Biochemically, GFAP consists of a central alpha-helical rod domain flanked by non-helical N-terminal head and C-terminal tail domains, responsible for filament assembly and stability (22). After TBI, GFAP and its breakdown products (*i.e.* GBDPs) are released into the blood stream though a complex interplay of BBB breakdown and glymphatic clearance (23). In human TBI, GFAP levels in blood are increased on ICU admission and are predictive of clinical outcomes at 6 months (24,25). The combination of blood GFAP and UCH-L1 biomarkers is FDA-approved for early detection of brain lesions after mild TBI (mTBI; 3,22), and GFAP alone predicts lesion positivity on neuroimaging scans in mTBI patients (9,27).

### Tau and p-Tau

Tau, a microtubule-associated protein found mainly in axons, plays a critical role in maintaining neuronal cytoskeletal structure. In its phosphorylated form (p-Tau), Tau regulates normal cellular function (28). However, aberrant post-translational modification, misfolding, and the assembly of Tau into filaments destabilizes microtubules, affects axonal transport, and disrupts neuronal function. p-Tau is implicated in the pathogenesis of Alzheimer’s Disease (AD) and other tauopathies, and it contributes to chronic neurodegeneration in TBI (29), including chronic traumatic encephalopathy (CTE) (30). p-Tau epitopes have been identified and validated in blood as biomarkers, with p-Tau231 and p-Tau217 plasma levels being strongly associated with early cerebral Aβ changes in AD patients (31). Plasma p-Tau:t-Tau ratio has shown higher diagnostic and prognostic accuracy over t-Tau levels for acute TBI and shows more robust and sustained elevations among patients with chronic TBI (32,33,34).

### UCH-L1

UCH-L1 is a 25 kDa cytoplasmic deubiquitinating enzyme primarily found in neuronal cell bodies where it is involved in axonal transport and regulates brain protein metabolism (35). Specifically, UCH-L1 regulates the ubiquitination of proteins marked for degradation via the ubiquitin-proteasome pathway. Dysfunction in the ubiquitin-proteasome pathway worsens secondary injury after TBI (36). Clinical investigations show an early increase in serum UCH-L1 in mild and moderate TBI patients detectable within 1-hour post-injury is associated with injury severity, CT lesions, and neurological intervention (35,37).

In the current study, we performed a systematic review of preclinical TBI studies in rodents, measuring blood protein biomarkers that are widely used in clinical studies. We focused on circulating protein biomarkers of TBI in blood, including NfL, t-Tau, p-Tau, UCH-L1, and GFAP (Fig.1). We discuss the kinetics of the preclinical biomarkers following varying degrees of injury severity and their potential to inform on secondary injury evolution and guide treatment strategies for TBI. Finally, we address how blood-based biomarker trajectories can be used to calibrate pathophysiological mechanisms in rodent TBI models with human disease, thus, enhancing bidirectional translational research in the field.

**Fig. 1.**
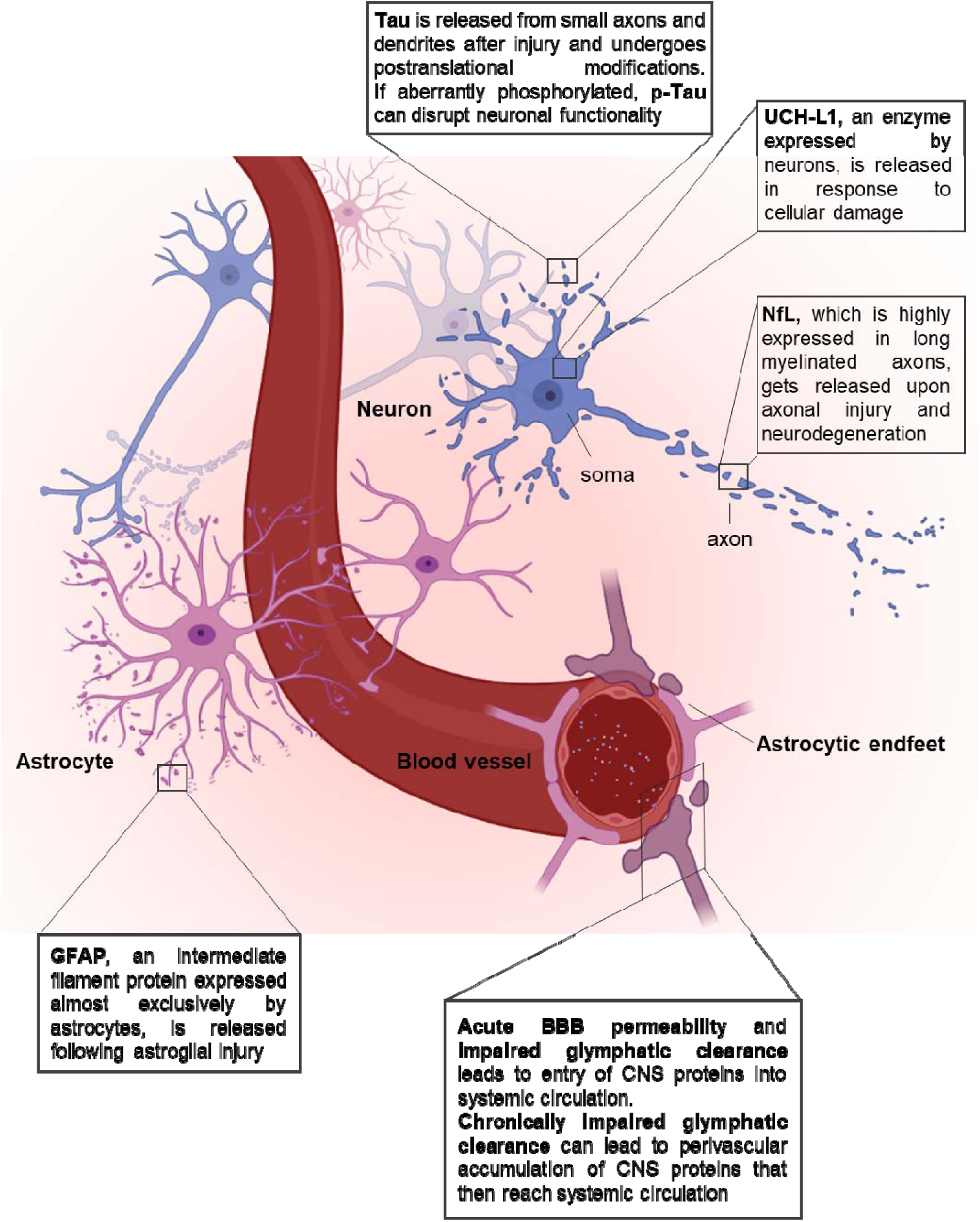
| Key protein biomarkers and pathobiological changes associated with TBI. Abbreviations: NfL: neurofilament-light; GFAP: glial fibrillary acidic protein; p-Tau: phosphorylated Tau; UCH-L1: ubiquitin C-terminal hydrolase; BBB: blood-brain barrier. Image generated using Biorender.

## Methods

This study was pre-registered on the International Prospective Register of Systematic Reviews (PROSPERO)—identification number: CRD42023423859 (https://www.crd.york.ac.uk/prospero/display_record.php?RecordID=423859).

## Search strategy and data extraction

PubMed was searched for English articles without date restriction up to 13^th^ June 2023. The search strategy used was: ((rat) OR (mouse) OR (rodent)) AND ((serum) OR (plasma) OR (blood)) AND ((traumatic brain injury) OR (TBI) OR (neurotrauma)) AND ((NFL) OR (NF-L) OR (Neurofilament) OR (TAU) OR (GFAP) OR (glial fibrillary acidic protein) OR (UCHL1) OR (UCH-L1)). We included all relevant preclinical rodent studies assessing at least one of the five main blood-biomarkers identified (NfL, Tau/ p-Tau, UCH-L1, GFAP). The screening of the titles, abstract and data extraction was performed by three independent reviewers (FM, IL & EM) and any disagreement was discussed and resolved by a fourth independent reviewer (FP). Duplicates, reviews, studies assessing other or non-blood-based biomarkers, and studies investigating biomarkers not in TBI were excluded. Data extraction sought to obtain information on the TBI model type and severity, age, sex and genetic background of the rodents used, the time points at which the biomarkers were measured, the main findings in relation to the biomarkers, the approximate biomarkers values with respect to controls, and the prognostic and pharmacodynamic potential of the marker. Whenever age was not reported, we used publicly available conversion tables to infer the rodents’ age from their weight. For ease of interpretation, the severities of injuries in each study were categorized into moderate-to-severe TBI, single mild (smTBI), or repetitive mild (rmTBI) based on the information provided in each study. When generating figures and interpreting the data, the time points across studies were grouped as follows: all blood samples collected during the initial 24 hours post-injury were designated as ‘hours’; between 24 hours and 7 days as ‘days’; between 7 days and 1 month as ‘weeks’; and any duration exceeding 1 month was labeled as ‘months’.

The quality of the studies included was assessed by two independent investigators (IL & EM), based on a checklist from the Collaborative Approach to Meta-Analysis and Review of Animal Data from Experimental Studies (CAMARADES) (23).

## Results

Our search for preclinical rodent studies investigating blood-based TBI biomarkers yielded 369 studies, of which 71 met eligibility criteria and were included for data extraction. A PRISMA flow diagram outlining each step of our search strategy is presented in **Fig. 2**.

**Fig. 2.**
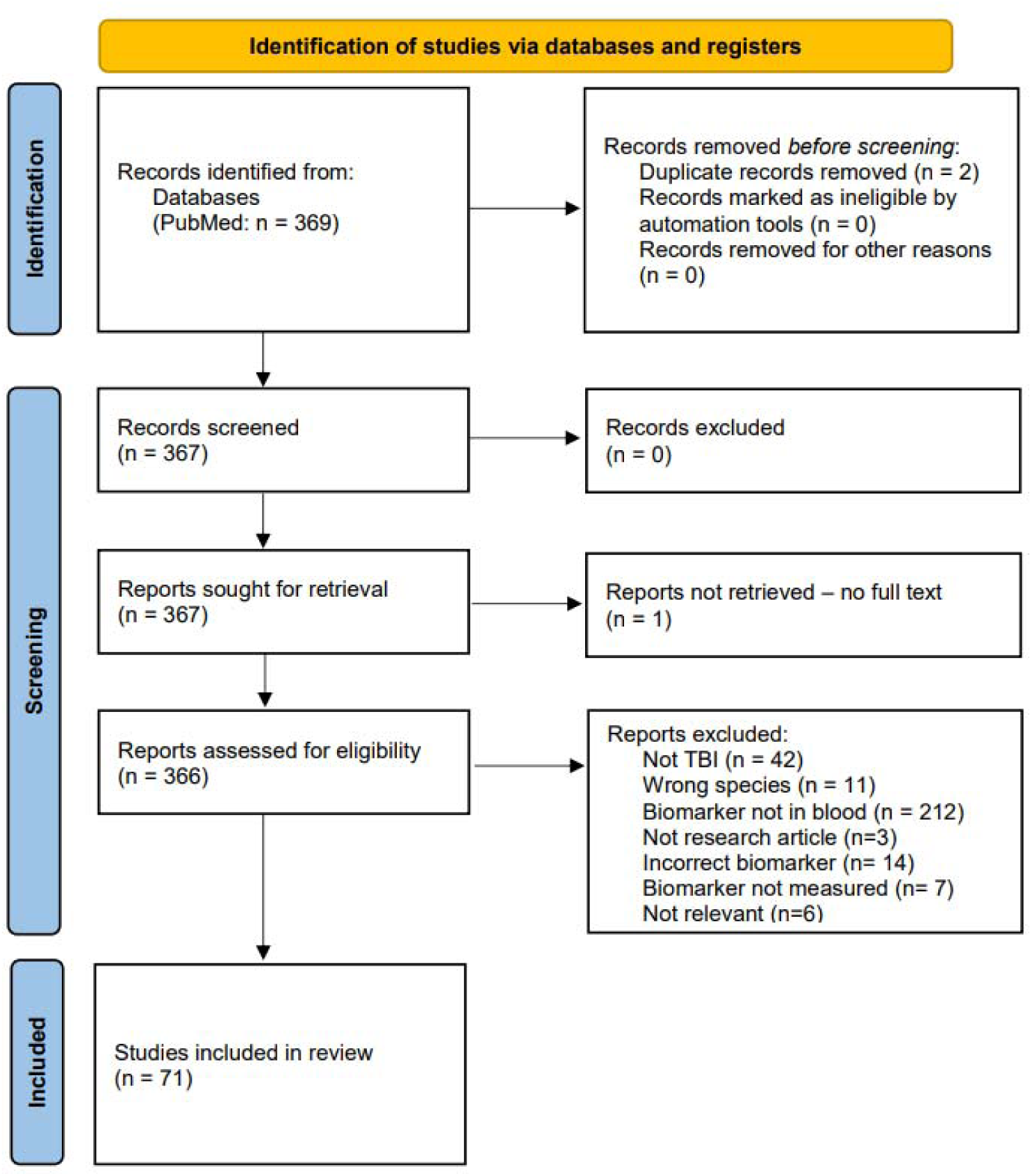
| PRISMA 2020 flow diagram for the systematic review.

Quality score was determined with the checklist modified from the CAMARADES (**Suppl. Table 1** for complete evaluation, **Fig. 3A-B** for summary data). The median quality score across the 71 studies was 5 (interquartile range 4–7). Seventeen studies assessed NfL, 41 GFAP, 17 t-Tau, 7 p-Tau, and 19 UCH-L1 (**Fig. 3C**), with 27 studies assessing more than one marker at a time. Thirty-seven studies assessed the biomarkers in moderate-to-severe TBI, 12 in smTBI and 9 in rmTBI. Five studies assessed both smTBI and moderate-to-severe injury, 7 smTBI and rmTBI, and 1 study assessed all three injury severities. Most studies investigated TBI in rats (n= 48), 22 in mice, and 1 in both species (**Fig. 3D**). Fifty-five studies assessed blood biomarkers in male animals, 8 in both males and females, 1 in female only, and 7 did not report the sex of the animals used (**Fig. 3E**). Sixty-five studies used adult rodents (of which for 23 studies age had to be inferred from the animal weight and was not otherwise reported), 2 pediatric, and 2 aged (and adult) rodents, while 2 did not report the age of the rodents used, nor their weight (**Fig. 3F**). Blood biomarkers were assessed in serum (n=48) or plasma (n=23). Seventeen studies investigated changes in the biomarkers in response to effective treatments (i.e., the biomarkers’ value as a pharmacodynamic biomarker). The techniques used for biomarkers’ measurement varied widely and are schematically represented in **Fig. 3G**, with ELISA being the most widely used technique to quantify protein biomarker levels (54%).

**Fig. 3.**
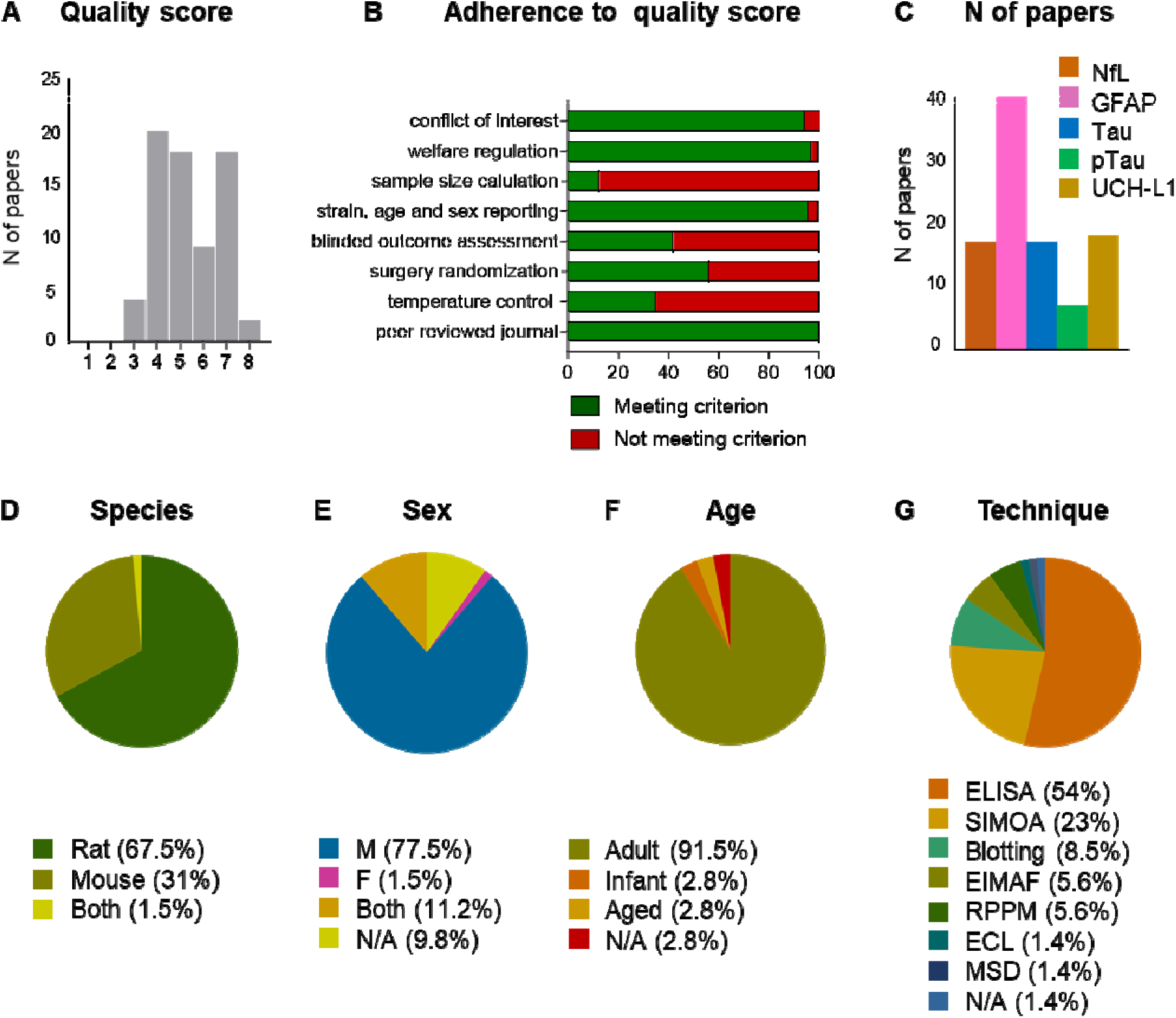
| Characteristics of the 71 studies included in the review. **A-B:** quality score and adherence to quality score criteria, as assessed by the CAMARADES checklist; **C:** number of papers investigating each biomarker; **D-G:** percentage of rodent species used, male or female, age group, and technique used for biomarker measurement. Abbreviations: NfL: neurofilament light; GFAP: glial fibrillary acidic protein; p-Tau: phosphorylated Tau; UCH-L1: ubiquitin C-terminal hydrolase; M: male; F: female; N/A: not available; ELISA: enzyme-linked immunoassay; SIMOA: single molecule array; EIMAF: Enhanced Immunoassay using Multi-Arrayed Fiberoptics; RPPM: reverse phase protein microarray; ECL: electrochemiluminescence; MSD: Meso Scale Discovery.

## NfL

Circulating blood NfL was assessed in 17 preclinical TBI studies (**Table 1**). Injury severity was moderate-to-severe TBI in 8 studies, smTBI in 5 studies, and rmTBI in 8 studies. Twelve studies performed a longitudinal assessment of NfL after injury (7,39–49).

**Table 1.** | NfL studies.

| Reference | Species | Sex | Genetic background | Age | Model | Severity | Biofluid | Assay format | Timepoints assessed | Pharmacodynamic assessment | Prognostic assessment |
| --- | --- | --- | --- | --- | --- | --- | --- | --- | --- | --- | --- |
| Bashir et al., 2020 | mouse | male and female | WT | adult | CHIMERA | moderate-to-severe | plasma | SIMOA | 6h | N | N |
| Boucher et al., 2022 | mouse | male | WT | adult | WD | rmTBI | serum | SIMOA | 1d, 1w, 1m | N | N |
| Cheng et al., 2018 | mouse | male | WT & TG (AAP/PS1) | adult and aged | CHIMERA | smTBI | plasma | SIMOA | 6h, 2d, 1w | N | N |
| Dickstein et al., 2021 | rat | male | WT | adult | BI | rmTBI | plasma | SIMOA | 4m, 10m | N | N |
| Diomedea et al., 2023 | mouse | male and female | WT | adult and aged | CCI | moderate-to-severe | plasma | SIMOA | 1w | Y | N |
| Graham et al., 2021 | rat | male | WT | adult | CCI | moderate-to-severe | plasma | SIMOA | 3d, 1w, 10d, 2w | N | Y |
| Heiskanen et al., 2022 | rat | male | WT | adult | FPI | moderate-to-severe | plasma | SIMOA | 2d, 9d, 6m | N | Y |
| Koza et al., 2023 | mouse | male | WT | adult | CHI | rmTBI | serum | ELISA | 3d | Y | N |
| Li et al., 2021 | rat | male | WT | adult | PBBI | moderate-to-severe | serum | SIMOA | 1d, 2d | N | N |
| Moro et al., 2022 | mouse | male and female | WT | adult | CHI | smTBI; rmTBI | plasma | SIMOA | 1w, 5w, 3m, 6m, 12m | N | Y |
| O'Brien et al., 2021 | rat | male | WT | adult | aCHI | smTBI; rmTBI | serum | SIMOA | 2h, 1d, 3d, 1w, 2w | N | N |
| O'Brien et al., 2023 | rat | male | WT | adult | aCHI | smTBI; rmTBI | serum | SIMOA | 3d, 1w, 2w, 1m | N | Y |
| Pham et al., 2021 | rat | male | WT | adult | aCHI | rmTBI | serum | SIMOA | 1w, 3.5m | N | N |
| Saletti et al., 2023 | rat | male | WT | adult | FPI | moderate-to-severe | plasma | RPPM | 2d, 1w | Y | Y |
| Stemper et al., 2022 | rat | male | WT | adult | HRA | smTBI; rmTBI | serum | SIMOA | 1w, 2w, 3w, 1m, 5w | N | N |
| Thau-Zuchman et al., 2020 | mouse | N/A | WT | adult | CCI | moderate-to-severe | plasma | ECL | 1m | Y | N |
| Wong et al., 2021 | rat | male | WT | adult | FPI | moderate-to-severe | serum | SIMOA | 2d | N | N |

Abbreviations: WT: wild-type; TG: transgenic; CHIMERA: closed-head impact model of engineered rotational acceleration; WD: weight drop; BI: blast injury; CCI: controlled cortical impact; FPI: fluid percussion injury; CHI: closed head injury; PBBI: penetrating ballistic brain injury; aCHI: awake closed head injury; HRA: head rotational acceleration; smTBI: single mild TBI; rmTBI: repetitive mild TBI; SIMOA: single molecule array; ELISA: enzyme-linked immunoassay; ECL: electrochemiluminescence; h: hours; d: days; w: weeks; m: months. N: no; Y: yes

### Biomarker kinetics

In moderate-to-severe TBI models, 2 studies assessed NfL levels within 0-24 hours (h) of injury (’hours’, (36,47), 6 studies from 1 to 6 days (d) (‘days’, (7,39–41,51,52)), 3 studies from 1 to 4 weeks (w) (‘weeks’, (7,40,53)), and only 1 study assessed NfL at months (m) post-injury (’months’, (37)) (**Fig. 4A**). Following moderate-to-severe TBI, NfL increased and peaked at 1-3d (7,29,30,42) with NfL levels remaining elevated up to 6m after TBI (40). In smTBI models, NfL levels were assessed at ‘hours’ by 2 studies (43,46), at ‘days’ by 5 (43,45–47,49), at ‘weeks’ by 4 studies (45–47,49), and at ‘months’ by 2 studies (45,49) (**Fig. 4B**). Following smTBI, NfL peaked between 6h (33) and 3d (46,47) post-injury, with NfL levels remaining elevated at 1w (45), 2w (46,47) and even 4w (49) after smTBI. NfL levels returned to baseline by 5w (45). In rmTBI models, NfL levels were assessed at ‘hours’ by 2 studies (42,46), at ‘days’ by 7 (42,45–49,54), at ‘weeks’ by 4 (42,46,47,49), and at ‘months’ by 4 studies (44,45,48,49) (**Fig. 4C**). Following rmTBI, there was a delayed peak in NfL levels between 3d (46) and 30d (49) post-injury. NfL levels were reported to return to baseline by 3m post-injury (45,47,48), with the exception of one study that showed elevated NfL levels at 4m following at single mild blast injury (44).

**Fig. 4.**
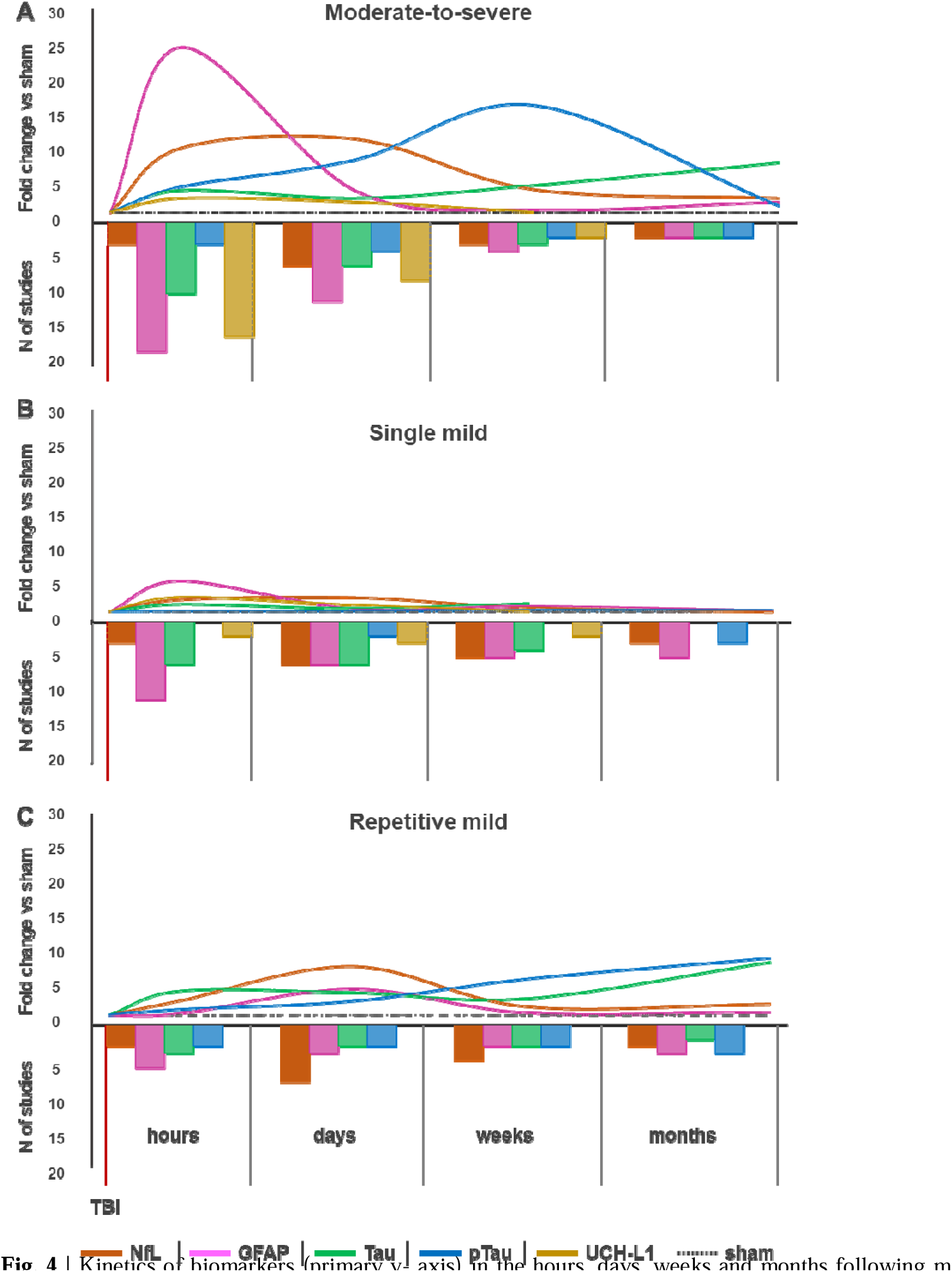
| Kinetics of biomarkers (primary y-axis) in the hours, days, weeks and months following moderate-to-severe injury. (**A**), smTBI (**B**) or rmTBI (**C**) TBI. On the secondary y axis are represented the number of studies assessing each biomarker at the timepoints of interest. Abbreviations: NfL: neurofilament-light; GFAP: glial fibrillary acidic protein; p-Tau: phosphorylated Tau; UCH-L1: ubiquitin C-terminal hydrolase

### Value as prognostic biomarker

The prognostic value of circulating NfL was assessed in four preclinical studies (40,41,45,46). Acute NfL levels were reported to be indicative of chronic lesion size after fluid percussion injury (40) and callosal atrophy (7,45), and to predict after rmTBI subacute neurological video signs of brain injury (including postimpact seizures, motionlessness and forelimb or hindlimb incoordination) (47). Two studies found that NfL did not predict chronic memory deficits as assessed using the Morris water maze test (40), nor sensorimotor function impairments as assessed using the composite neuroscore (41).

### Value as a pharmacodynamic biomarker

The pharmacodynamic value of circulating NfL was assessed in four preclinical drug treatment studies (41,51,53,54). Drugs tested included a) Aβ1-6A2V(D), an all-D-isomer synthetic peptide that interferes with the aggregation and stability of tau protein (51); b) Immunocal, a cysteine-rich whey protein supplement and glutathione precursor (54); c) levetiracetam (LEV), an anti-seizure medication (41); and d) docosahexaenoic acid (DHA), a neuroreactive omega-3 fatty acid (53). All drugs had demonstrated neurotherapeutic effects in preclinical studies based on other neurological, biochemical, and histological outcome measures. Of these, Aβ1-6A2V(D) and DHA reported a decrease TBI-induced blood NfL levels following drug treatment (51,53), while the other two drugs (Immunocal, LEV) did not (41,54).

## GFAP

Circulating blood GFAP levels were assessed in 41 preclinical TBI studies (**Table 2**). Injury severity was moderate-to-severe TBI in 26 studies, smTBI in 15, and rmTBI in 7 studies. Twenty-six studies performed a longitudinal assessment of GFAP levels after TBI (41,42,55–80).

**Table 2.**
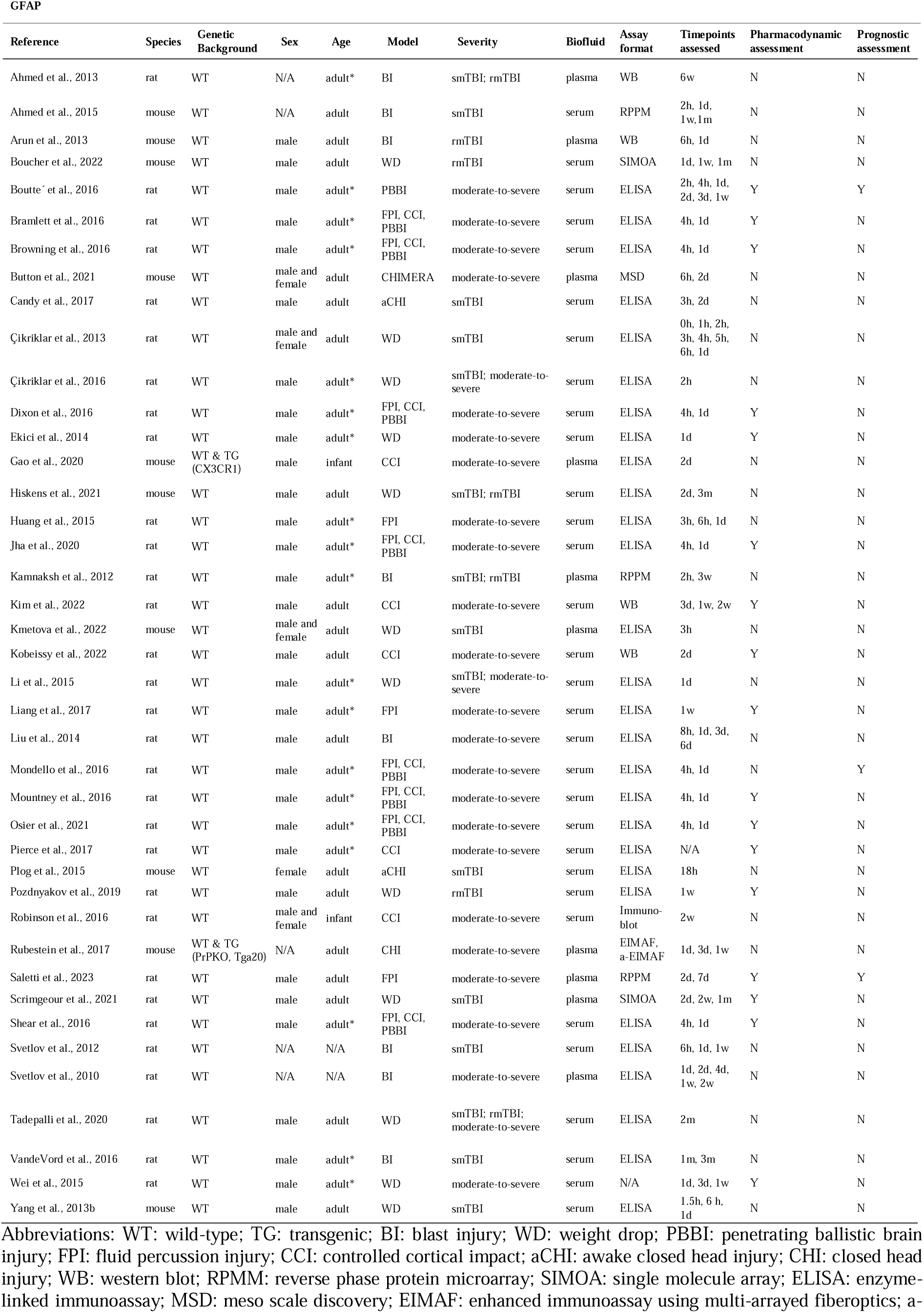
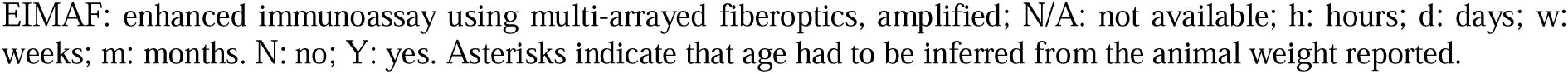
| GFAP studies.

| GFAP |  |  |  |  |  |  |  |  |  |  |  |
| --- | --- | --- | --- | --- | --- | --- | --- | --- | --- | --- | --- |
| Reference | Species | Genetic Background | Sex | Age | Model | Severity | Biofluid | Assay format | Timepoints assessed | Pharmacodynamic assessment | Prognostic assessment |
| Ahmed et al., 2013 | rat | WT | N/A | adult* | BI | smTBI; rmTBI | plasma | WB | 6w | N | N |
| Ahmed et al., 2015 | mouse | WT | N/A | adult | BI | smTBI | serum | RPPM | 2h, 1d, 1w, 1m | N | N |
| Arun et al., 2013 | mouse | WT | male | adult | BI | rmTBI | plasma | WB | 6h, 1d | N | N |
| Boucher et al., 2022 | mouse | WT | male | adult | WD | rmTBI | serum | SIMOA | 1d, 1w, 1m | N | N |
| Boutte<sup>c</sup> et al., 2016 | rat | WT | male | adult* | PBBI | moderate-to-severe | serum | ELISA | 2h, 4h, 1d, 2d, 3d, 1w | Y | Y |
| Bramlett et al., 2016 | rat | WT | male | adult* | FPI, CCI, PBBI | moderate-to-severe | serum | ELISA | 4h, 1d | Y | N |
| Browning et al., 2016 | rat | WT | male | adult* | FPI, CCI, PBBI | moderate-to-severe | serum | ELISA | 4h, 1d | Y | N |
| Button et al., 2021 | mouse | WT | male and female | adult | CHIMERA | moderate-to-severe | plasma | MSD | 6h, 2d | N | N |
| Candy et al., 2017 | rat | WT | male | adult | aCHI | smTBI | serum | ELISA | 3h, 2d | N | N |
| Çikriklar et al., 2013 | rat | WT | male and female | adult | WD | smTBI | serum | ELISA | 0h, 1h, 2h, 3h, 4h, 5h, 6h, 1d | N | N |
| Çikriklar et al., 2016 | rat | WT | male | adult* | WD | smTBI; moderate-to-severe | serum | ELISA | 2h | N | N |
| Dixon et al., 2016 | rat | WT | male | adult* | FPI, CCI, PBBI | moderate-to-severe | serum | ELISA | 4h, 1d | Y | N |
| Ekici et al., 2014 | rat | WT | male | adult* | WD | moderate-to-severe | serum | ELISA | 1d | Y | N |
| Gao et al., 2020 | mouse | WT & TG (CX3CR1) | male | infant | CCI | moderate-to-severe | plasma | ELISA | 2d | N | N |
| Hiskens et al., 2021 | mouse | WT | male | adult | WD | smTBI; rmTBI | serum | ELISA | 2d, 3m | N | N |
| Huang et al., 2015 | rat | WT | male | adult* | FPI | moderate-to-severe | serum | ELISA | 3h, 6h, 1d | N | N |
| Jha et al., 2020 | rat | WT | male | adult* | FPI, CCI, PBBI | moderate-to-severe | serum | ELISA | 4h, 1d | Y | N |
| Kamnakhsh et al., 2012 | rat | WT | male | adult* | BI | smTBI; rmTBI | plasma | RPPM | 2h, 3w | N | N |
| Kim et al., 2022 | rat | WT | male | adult | CCI | moderate-to-severe | serum | WB | 3d, 1w, 2w | Y | N |
| Kmetova et al., 2022 | mouse | WT | male and female | adult | WD | smTBI | plasma | ELISA | 3h | N | N |
| Kobeissy et al., 2022 | rat | WT | male | adult | CCI | moderate-to-severe | serum | WB | 2d | Y | N |
| Li et al., 2015 | rat | WT | male | adult* | WD | smTBI; moderate-to-severe | serum | ELISA | 1d | N | N |
| Liang et al., 2017 | rat | WT | male | adult* | FPI | moderate-to-severe | serum | ELISA | 1w | Y | N |
| Liu et al., 2014 | rat | WT | male | adult | BI | moderate-to-severe | serum | ELISA | 8h, 1d, 3d, 6d | N | N |
| Mondello et al., 2016 | rat | WT | male | adult* | FPI, CCI, PBBI | moderate-to-severe | serum | ELISA | 4h, 1d | N | Y |
| Mountney et al., 2016 | rat | WT | male | adult* | FPI, CCI, PBBI | moderate-to-severe | serum | ELISA | 4h, 1d | Y | N |
| Osier et al., 2021 | rat | WT | male | adult* | FPI, CCI, PBBI | moderate-to-severe | serum | ELISA | 4h, 1d | Y | N |
| Pierce et al., 2017 | rat | WT | male | adult* | CCI | moderate-to-severe | serum | ELISA | N/A | Y | N |
| Plog et al., 2015 | mouse | WT | female | adult | aCHI | smTBI | serum | ELISA | 18h | N | N |
| Pozdnyakov et al., 2019 | rat | WT | male | adult | WD | rmTBI | serum | ELISA | 1w | Y | N |
| Robinson et al., 2016 | rat | WT | male and female | infant | CCI | moderate-to-severe | serum | Immuno-blot | 2w | N | N |
| Rubenstein et al., 2017 | mouse | WT & TG (PrPKO, Tga20) | N/A | adult | CHI | moderate-to-severe | plasma | EIMAF, a-EIMAF | 1d, 3d, 1w | N | N |
| Saletti et al., 2023 | rat | WT | male | adult | FPI | moderate-to-severe | plasma | RPPM | 2d, 7d | Y | Y |
| Scrimgeour et al., 2021 | rat | WT | male | adult | WD | smTBI | plasma | SIMOA | 2d, 2w, 1m | Y | N |
| Shear et al., 2016 | rat | WT | male | adult* | FPI, CCI, PBBI | moderate-to-severe | serum | ELISA | 4h, 1d | Y | N |
| Svetlov et al., 2012 | rat | WT | N/A | N/A | BI | smTBI | serum | ELISA | 6h, 1d, 1w | N | N |
| Svetlov et al., 2010 | rat | WT | N/A | N/A | BI | moderate-to-severe | plasma | ELISA | 1d, 2d, 4d, 1w, 2w | N | N |
| Tadepalli et al., 2020 | rat | WT | male | adult | WD | smTBI; rmTBI; moderate-to-severe | serum | ELISA | 2m | N | N |
| VandeVord et al., 2016 | rat | WT | male | adult* | BI | smTBI | serum | ELISA | 1m, 3m | N | N |
| Wei et al., 2015 | rat | WT | male | adult* | WD | moderate-to-severe | serum | N/A | 1d, 3d, 1w | Y | N |
| Yang et al., 2013b | mouse | WT | male | adult | WD | smTBI | serum | ELISA | 1.5h, 6 h, 1d | N | N |

Abbreviations: WT: wild-type; TG: transgenic; BI: blast injury; WD: weight drop; PBBI: penetrating ballistic brain injury; FPI: fluid percussion injury; CCI: controlled cortical impact; aCHI: awake closed head injury; CHI: closed head injury; WB: western blot; RPPM: reverse phase protein microarray; SIMOA: single molecule array; ELISA: enzyme-linked immunoassay; MSD: meso scale discovery; EIMAF: enhanced immunoassay using multi-arrayed fiberoptics; a-
EIMAF: enhanced immunoassay using multi-arrayed fiberoptics, amplified; N/A: not available; h: hours; d: days; w: weeks; m: months. N: no; Y: yes. Asterisks indicate that age had to be inferred from the animal weight reported.

### Biomarker kinetics

In moderate-to-severe TBI models, circulating GFAP levels were assessed at ‘hours’ in 18 studies (57–59,62,64,65,68–72,74,75,78,80–83), at ‘days’ in 11 studies (41,57,67,68,72,75,78,80,84–86), ‘weeks’ in 3 studies (67,75,87), and ‘months’ in 1 study (88) (**Fig. 4A**). One study did not report the timepoint of GFAP sampling (89). Following moderate-to-severe TBI, GFAP sharply increased within 2h post-injury (57,80), peaked at 4h-24h post-injury (57–59,64,69–71,74), and returned to sham levels at one week post-injury (41,57,75,80). Only one study reported GFAP levels increased at 2 months post-injury (88). In smTBI models, GFAP was assessed at ‘hours’ by 10 studies (55,60,61,66,76,79,81,83,90,91), at ‘days’ by 5 studies (55,60,63,73,76), at ‘weeks’ by 4 studies (55,66,73,77), and at ‘months’ by 4 studies (63,77,88,92) (**Fig. 4B**). Following smTBI, GFAP was reported to marginally increase in the hours after smTBI (61,75,79,83,90), and then return to sham values between 2-7d post-injury (56,60,63). There was an exception in one study using a weight drop injury model that reported increased GFAP levels at 30d post-injury (73). In rmTBI models, GFAP was assessed at ‘hours’ in 5 studies (42,56,66,81,83), at ‘days’ in 3 studies (42,63,93), at ‘weeks’ in 2 studies (42,66), and at ‘months’ in 3 studies (63,88,92) (**Fig. 4C**). Following rmTBI, there was a slower increase in blood GFAP levels detected at 1d post-injury (56), and increased GFAP levels were observed many weeks after rmTBI (66,88,92,93).

### Value as prognostic biomarker

The prognostic value of circulating GFAP was assessed in 3 preclinical studies (41,57,69). Blood GFAP levels at 4h post-injury predicted motor function impairment, assessed through the composite neuroscore (57) and contusion volume/tissue loss at three weeks post-injury (69). Blood GFAP levels at 24h also correlated with tissue GBDPs levels on day 3 post-injury (57), and were predictive of contusion volume/tissue loss at week 3 post-injury (69). One study reported no prognostic value of GFAP after TBI (41).

### Value as a pharmacodynamic biomarker

The pharmacodynamic value of GFAP was assessed in 17 preclinical studies (41,57–59,62,65,67,70,71,73,74,78,82,85,86,89,93) using various injury severity models. Different therapeutic approaches were evaluated and ranged from lifestyle interventions (67,73,86), to anti-epileptic drugs, (41,59) to immunomodulatory agents (57,62,82). TBI-induced blood GFAP levels were decreased in 9 out of 13 studies from moderate-to-severe TBI (57,59,67,74,78,85,89), smTBI (73), and rmTBI (93) models.

## t-Tau and p-Tau

Circulating blood t-Tau was assessed in 20 preclinical TBI studies and blood p-Tau was assessed in 9 preclinical TBI studies (**Table 3**). For blood t-Tau, injury severity was moderate-to-severe TBI in 10 studies, smTBI in 6 studies, and rmTBI in 4 studies. For blood p-Tau, injury severity was moderate-to-severe TBI in 4 studies, smTBI in 1 study, and rmTBI in 4 studies. Fourteen studies performed a longitudinal assessment of t-Tau and p-Tau after TBI (41,43,55,56,63,68,72,73,94–99).

**Table 3.** | t-Tau and p-Tau studies.

| Reference | Species | Genetic Background | Sex | Age | Model | Severity | Biofluid | Assay format | Timepoints assessed | Pharmacodynamic assessment | Prognostic assessment |
| --- | --- | --- | --- | --- | --- | --- | --- | --- | --- | --- | --- |
| Ahmed et al., 2015 | mouse | WT | N/A | adult | BI | smTBI | serum | RPPM | 2h, 1d, 1w, 1m | N | N |
| Arun et al., 2013 | mouse | WT | male | adult | BI | rmTBI | plasma | WB | 6h, 1d | N | N |
| Bashir et al., 2020 | mouse | WT | male and female | adult | CHIMERA | moderate-to-severe | plasma | SIMOA | 6h | N | N |
| Cheng et al., 2023 | mouse | TG (rTg4510) | male | adult | CHIMERA | moderate-to-severe | plasma | SIMOA | 6h, 2w, 1m, 1.5m, 2m | N | N |
| Ekici et al., 2014 | rat | WT | male | adult* | WD | moderate-to-severe | serum | ELISA | 1d | Y | N |
| Li et al., 2020 | rat | WT | male | adult | BI | smTBI | serum | ELISA | 1d, 3d | Y | N |
| Liliang et al., 2010 | rat | WT | male | adult* | WD | smTBI; moderate-to-severe | serum | ELISA | 1h, 6h, 1d, 2d, 1w | N | N |
| Liu et al., 2014 | rat | WT | male | adult | BI | moderate-to-severe | serum | ELISA | 8h, 1d, 3d, 6d | N | N |
| Rostami et al., 2012 | rat | WT | male | adult | WD | smTBI | serum | RPPM | 1d, 3d, 2w | N | N |
| Rubenstein et al., 2019 | mouse | TG (TghTau/PS) | male and female | adult | CHI | rmTBI | plasma | EIMAF | 1d, 10d, 2m, 4m, 6m, 8m, 10m, 12m | N | N |
| Rubenstein et al., 2015 | rat and mouse | WT | male | adult | CCI | moderate-to-severe | serum | EIMAF, a-EIMAF | 2h, 1d, 2d, 3d, 1w, 2w, 1m | N | N |
| Rubenstein et al., 2017 | mouse | WT & TG (PrPKO, Tga20) | male and female | adult | CHI | moderate-to-severe | plasma | EIMAF, a-EIMAF | 1d, 3d, 1w | N | N |
| Saletti et al., 2023 | rat | WT | male | adult | FPI | moderate-to-severe | plasma | RPPM | 2d, 7d | Y | Y |
| Scrimgeour et al., 2021 | rat | WT | male | adult | WD | smTBI | plasma | SIMOA | 2d, 2w, 1m | Y | N |
| Siahaan et al., 2018 | rat | WT | N/A | adult | WD | rmTBI | plasma | ELISA | 1w | Y | N |
| Tomita et al., 2020 | rat | WT | male | adult* | WD | smTBI; moderate-to-severe | serum | ELISA | 1h | N | N |
| Yang et al., 2015 | mouse | WT | male | adult | CHI | rmTBI | serum | EIMAF, a-EIMAF | 1d, 3d, 1w, 10d, 2w, 1m | N | N |
| Yao et al 2008 | rat | WT | male | adult* | PBBI | moderate-to-severe | plasma | WB | 1d | N | N |

**p-Tau**
| Reference | Species | Genetic Background | Sex | Age | Model | Severity | Biofluid | Assay format | Timepoints assessed | Pharmacodynamic assessment | Prognostic assessment |
| --- | --- | --- | --- | --- | --- | --- | --- | --- | --- | --- | --- |
| Hiskens et al., 2021 | mouse | WT | male | adult | WD | smTBI; rmTBI | serum | ELISA | 2d, 3m | N | N |
| Rubenstein et al., 2019 | mouse | TG (TghTau/PS) | male and female | adult | CHI | rmTBI | plasma | EIMAF | 1d, 10d, 2m, 4m, 6m, 8m, 10m, 12m | N | N |
| Rubenstein et al., 2015 | rat and mouse | WT | male | adult | CCI | moderate-to-severe | serum | EIMAF, a-EIMAF | 2h, 1d, 2d, 3d, 1w, 2w, 1m | N | N |
| Rubenstein et al., 2017 | mouse | WT & TG (PrPKO, Tga20) | male and female | adult | CHI | moderate-to-severe | plasma | EIMAF, a-EIMAF | 1d, 3d, 1w | N | N |
| Saletti et al., 2023 | rat | WT | male | adult | FPI | moderate-to-severe | plasma | RPPM | 2d, 7d | Y | Y |
| Tadepalli et al., 2020 | rat | WT | male | adult | WD | smTBI; rmTBI; moderate-to-severe | serum | ELISA | 2m | N | N |
| Yang et al., 2015 | mouse | WT | male | adult | CHI | rmTBI | serum | EIMAF, a-EIMAF | 1d, 3d, 1w, 10d, 2w, 1m | N | N |

Abbreviations: WT: wild-type; TG: transgenic; BI: blast injury; CHIMERA: closed-head impact model of engineered rotational acceleration; WD: weight drop; CHI: closed head injury; CCI: controlled cortical impact; PBBI: penetrating ballistic brain injury; RPPM: reverse phase protein microarray; WB: western blot; SIMOA: single molecule array; ELISA: enzyme-linked immunoassay; EIMAF: enhanced immunoassay using multi-arrayed fiber optics; a-EIMAF: enhanced immunoassay using multi-arrayed fiber optics, amplified; h: hours; d: days; w: weeks; m: months. N: no; Y: yes. \*: Asterisks indicate that age had to be inferred from the animal weight reported.

### Biomarker kinetics

In moderate-to-severe TBI models, circulating t-Tau levels were assessed at ‘hours’ in 9 studies (50,68,72,82,95,98,100–102) and circulating pTau levels in 2 studies (72,98). t-Tau was assessed at ‘days’ in 5 studies (41,68,72,95,98) and p-Tau in 3 studies (41,72,98). t-Tau was assessed at ‘weeks’ in 2 studies (98,100) and p-Tau in 1 study (98). t-Tau was assessed at ‘months’ in 1 study (100) and p-Tau in 1 study (88) (**Fig. 4A**). Following moderate-to-severe TBI, there was an acute increase in t-Tau within hours of TBI (68,95,101), and these levels remained elevated at weeks post-injury (98,100). For p-Tau, there was a slower increase in p-Tau starting within days of TBI (72,98), with a steady increase in p-Tau in the weeks (98) and months (88) post-injury. In smTBI models, t-Tau was assessed at ‘hours’ in 5 studies (55,94–96,101) and none for p-Tau. t-Tau was assessed at ‘days’ in 5 studies (55,73,94–96) and p-Tau in 1 study (63). t-Tau was assessed at ‘weeks’ in 3 studies (55,73,96) and none for p-Tau. There were no t-Tau studies at ‘months’, while pTau was assessed in 2 studies (63,88) (**Fig. 4B**). Following smTBI, there was an increase in t-Tau 1-6h after injury (55,95,101), with values elevated compared to sham at 30d post-injury (73). There were no changes in p-Tau levels compared to sham after smTBI (63,88). For rmTBI models, t-Tau was assessed ‘at hours’ in 3 studies (56,97,99) and p-Tau in 2 studies (97,99). t-Tau was assessed at ‘days’ in 2 studies (99,103) and p-Tau in 2 studies (63,99). t-Tau was assessed at ‘weeks’ in 2 studies (97,99) and p-Tau in 2 studies (97,99). t-Tau was assessed at ‘months’ in 1 study (97) and p-Tau in 3 studies (63,88,97) (**Fig. 4C**). Following rmTBI, both t-Tau and p-Tau gradually increase over time, from 24h up to 14d,(56,99), with values persistently elevated up to 1-year post-injury (97).

### Value as prognostic biomarker

Only one preclinical TBI study assessed the prognostic value of circulating t-Tau and p-Tau levels, reporting that both biomarkers predicted the occurrence of post-traumatic seizures (41).

### Value as a pharmacodynamic biomarker

The pharmacodynamic value of t-Tau was assessed in 5 preclinical studies (41,73,82,94,103), and in one study for p-Tau (41). Notably, TBI-induced circulating t-Tau and p-Tau levels were decreased following early normobaric hyperoxia and hyperbaric oxygen treatment (94), turmeric supplementation (103), and supplementation with omega-3 fatty acids and vitamin D3 (73). However, two studies reported no changes in circulating t-Tau and p-Tau in spite of reported neuroprotective effects of etanercept and lithium chloride (82) and LEV (41) in the experimental TBI models.

## UCH-L1

Circulating blood UCH-L1 was assessed in 20 preclinical TBI studies (**Table 4**). Injury severity was moderate-to-severe TBI in 18 studies, and smTBI in 2 studies. Fifteen studies performed a longitudinal assessment of UCH-L1 (57–59,62,64,65,70,71,73–75,104–107).

**Table 4.** | UCH-L1 studies.

| Reference | Species | Genetic Background | Sex | Age | Model | Severity | Biofluid | Assay format | Timepoints assessed | Pharmacodynamic assessment | Prognostic assessment |
| --- | --- | --- | --- | --- | --- | --- | --- | --- | --- | --- | --- |
| Aleem et al., 2022 | mouse | WT | male | adult | FPI | smTBI; moderate-to-severe | serum | ELISA | 1d, 3d | N | N |
| Boutte' et al., 2016 | rat | WT | male | adult* | PBBI | moderate-to-severe | serum | ELISA | 2h, 4h, 1d, 2d, 3d, 1w | N | N |
| Bramlett et al., 2016 | rat | WT | male | adult* | FPI, CCI, PBBI | moderate-to-severe | serum | ELISA | 4h, 1d | Y | N |
| Browning et al., 2016 | rat | WT | male | adult* | FPI, CCI, PBBI | moderate-to-severe | serum | ELISA | 4h, 1d | Y | N |
| Dixon et al., 2016 | rat | WT | male | adult* | FPI, CCI, PBBI | moderate-to-severe | serum | ELISA | 4h, 1d | Y | N |
| Huang et al., 2015 | rat | WT | male | adult* | FPI | moderate-to-severe | serum | ELISA | 3h, 6h, 1d | N | N |
| Jha et al., 2020 | rat | WT | male | adult* | FPI, CCI, PBBI | moderate-to-severe | serum | ELISA | 4h, 1d | Y | N |
| Kobeissy et al., 2022 | rat | WT | male | adult | CCI | moderate-to-severe | serum | WB | 2d | Y | N |
| Liu et al., 2010 | rat | WT | male | adult | CCI | moderate-to-severe | serum | ELISA | 2h, 6h, 12h, 1d, 1.5d, 2d, 2.5d, 3d | N | N |
| Mondello et al., 2016 | rat | WT | male | adult* | FPI, CCI, PBBI | moderate-to-severe | serum | ELISA | 4h, 1d | N | N |
| Morris et al., 2019 | mouse | WT | male | adult | WD | moderate-to-severe | serum | ELISA | 4h | N | N |
| Mountney et al., 2016 | rat | WT | male | adult* | FPI, CCI, PBBI | moderate-to-severe | serum | ELISA | 4h, 1d | Y | N |
| Osier et al., 2021 | rat | WT | male | adult* | FPI, CCI, PBBI | moderate-to-severe | serum | ELISA | 4h, 1d | Y | N |
| Pierce et al., 2017 | rat | WT | male | adult* | CCI | moderate-to-severe | serum | ELISA | N/A | Y | N |
| Scrimgeour et al., 2021 | rat | WT | male | adult | WD | smTBI | plasma | SIMOA | 2d, 2w, 1m | Y | N |
| Shear et al., 2016 | rat | WT | male | adult* | FPI, CCI, PBBI | moderate-to-severe | serum | ELISA | 4h, 1d | Y | N |
| Svetlov et al., 2010 | rat | WT | N/A | N/A | BI | moderate-to-severe | plasma | ELISA | 1d, 2d, 4d, 1w, 2w | N | N |
| Wallen et al., 2022 | mouse | WT | male | adult | WD | moderate-to-severe | serum | ELISA | 1h, 4h, 1d | N | N |
| Zoltewicz et al., 2013 | rat | WT | male | adult* | PBBI | moderate-to-severe | plasma | ELISA | 5min, 2h, 6h, 1d, 3d, 1w | N | N |

Abbreviations: WT: wild-type; TG: transgenic; FPI: fluid percussion injury; PBBI: penetrating ballistic brain injury; CCI: controlled cortical impact; WD: weight drop; BI: blast injury; ELISA: enzyme-linked immunoassay; WB: western blot; SIMOA: single molecule array; h: hours; d: days; w: weeks; m: months; N: no; Y: yes. Asterisks indicate that age had to be inferred from the animal weight reported.

### Biomarker kinetics

In moderate-to-severe TBI models, circulating UCH-L1 was assessed at ‘hours’ in 15 studies (57–59,62,64,65,70,71,74,75,104–108), at ‘days’ in 7 studies (57,70,73,75,85,104,105), and at ‘weeks’ in 1 study (73) (**Fig. 4A**). No studies assessed changes in UCH-L1 in the ‘months’ post-injury. One study (89) did not report the timepoint that UCH-L1 was assessed. Following moderate-to-severe TBI, there was a minor increase (2-3x sham values) in circulating UCH-L1 levels between 4h and 24h (57,58,62,64,65,71,75,104–107), and UCH-L1 levels returned back to sham levels by 24h (65,70,74) (**Fig. 4A**). There were two exceptions and both studies reported increased circulating UCH-L1 levels up to 3d (104) and 7d after injury (57). In smTBI models, circulating UCH-L1 was assessed at ‘hours’ in 1 study (104), at ‘days’ by 2 studies (73,104), and at ‘weeks’ by 1 study (73) (**Fig. 4B**). Following smTBI, UCH-L1 was modestly increased up to 3d post-injury (104), with one study reporting increased UCH-L1 levels at 30d post-injury, but not earlier (73). Circulating UCH-L1 biomarkers have not been assessed in rodent rmTBI.

### Value as prognostic biomarker

The prognostic value of circulating UCH-L1 levels has not been evaluated in rodent TBI models.

### Value as pharmacodynamic biomarker

The pharmacodynamic value of circulating UCH-L1 was assessed in 10 preclinical drug treatment studies (58,59,62,65,70,71,73,74,85,89). TBI-induced circulating UCH-L1 levels were decreased following treatment with LEV (59), ubiquinol (89), clopidogrel and aspirin (85), and dietary supplementation with omega-3 fatty acids and vitamin D3 (73). This was not the case for treatment with the membrane resealing agent Kollidon VA64 (71), Glibenclamide (65), nor Nicotinamide (74), all of which had significant neuroprotective effects as assessed using other measures. Three studies reported no therapeutic benefit of Cyclosporine-A (62), Erythropoietin (58), or Simvastatin (70) treatment, and no changes in circulating post-traumatic UCH-L1 levels.

## Discussion

This systematic literature review highlights that all blood-based biomarkers are increased after experimental TBI and show injury-severity dependency. GFAP and UCH-L1 peak in the hours after injury, while NfL shows a persistent increase in the days following TBI. There is a delayed increase in t-Tau and p-Tau in the weeks following TBI. The available data in rmTBI models suggest that blood biomarker level peaks are delayed, pointing to a sensitivity to repeated brain injury. Since the pattern of biomarker changes mimic what is observed in human TBI (3,14,35), we next discuss the main findings for each biomarker and, whenever possible, draw connections with existing knowledge from human TBI studies and/or gyroencephalic preclinical models.

NfL is highly sensitive to TBI in preclinical models, with only three studies reporting no variations in NfL after either moderate-to-severe (41), smTBI (27) or rmTBI (28). Across multiple models of single TBI, including closed head injury, controlled cortical impact, penetrating injury, blast injury and CHIMERA, NfL peaks in the days following injury (**Fig. 4A-C**) (7,39,40,46,50,52), with an injury-severity-dependent elevation (7) indicating that axonal injury/damage is a common pathophysiological mechanism. Notably, following rmTBI, NfL responds to cumulative injuries (45,46,49) and injury intervals (45,47), suggesting that NfL levels could inform on susceptibility to subsequent mild TBIs. The overall temporal pattern of NfL blood levels closely aligns with studies in gyroencephalic models in pigs (109), and with the human data (7,20). However, there is a difference in the time-to-first-peak between rodents (days 1-3, (39,40,52)), pigs (days 5-10,(109)), and humans (days 14-21; (20)) (**Fig. 5**), possibly related to species-specific differences in protein turnover rates or clearance mechanisms (110), anatomical differences in white matter (111), and differences in BBB dysfunction or permeability (112). The fact that circulating NfL levels in rodent models peak within days post-injury and remain elevated for about 1-2 weeks (depending on injury severity) is a significant advantage for preclinical research and can be incorporated into experimental design to reduce sample size requirements and decrease experimental group variability. It is important to note that peripheral nerve damage (113) and physiological aging (114), can also increase NfL levels. Studies in rodents (52) and patients (6) suggest that following severe TBI the contribution of injured peripheral nerves to NfL blood levels is negligible. Nonetheless, how peripheral injuries influence circulating NfL in mTBI with co-occurring systemic injuries has not been examined to date. Therefore, preclinical models represent the ideal setting to dissect the specific contribution of central and peripheral injuries to circulating NfL levels following mTBI.

**Fig. 5.**
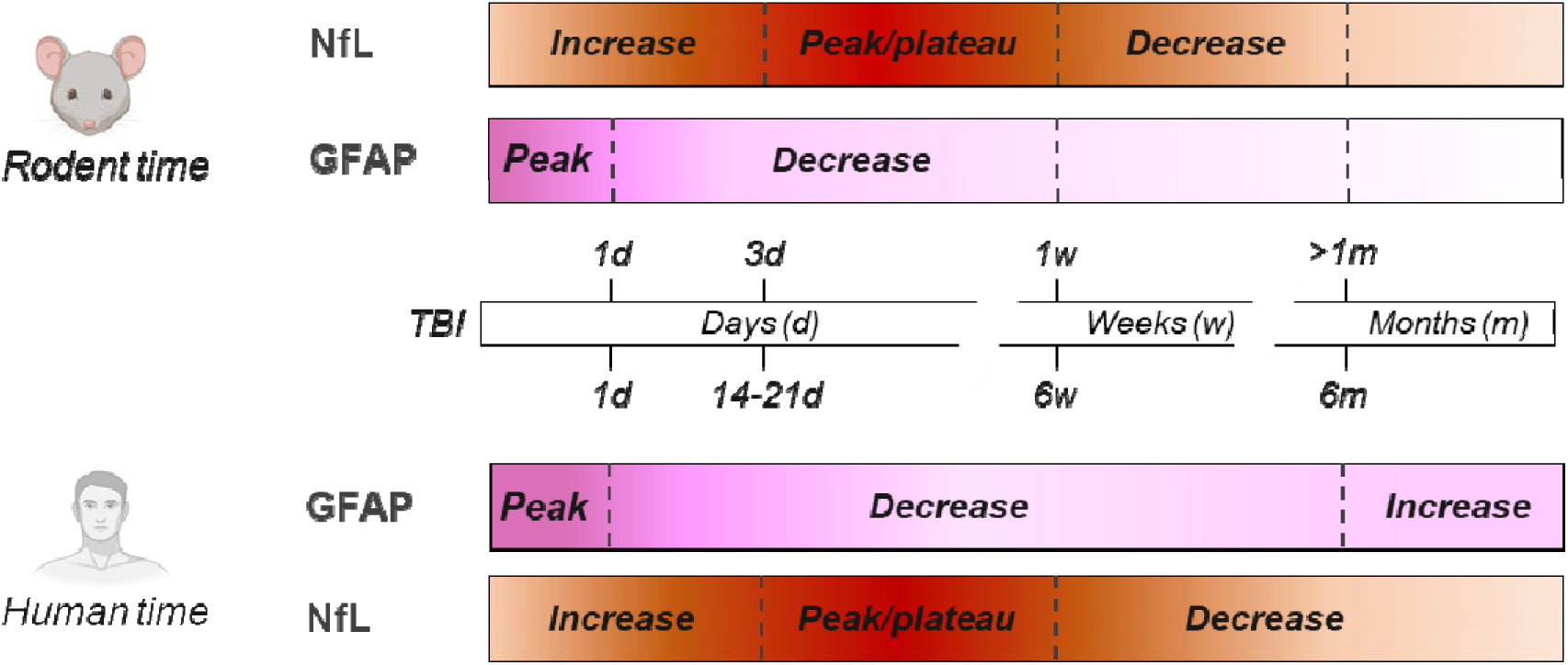
| NfL and GFAP temporal dynamics after moderate-to-severe TBI in rodents (above timeline) and humans (below timeline)

Circulating NfL levels correlate with sensorimotor (hyperactivity in the open field test, 30; beam walk, 31) and cognitive (Morris water maze, 25) impairments and predicts evolution of structural damage, including lesion size (40) and callosal atrophy (45) following TBI. These findings align with observations in injured pigs (109) and humans (7,20), highlighting the cross-species prognostic value of post-traumatic NfL levels in predicting brain volume loss and white matter abnormalities. In addition, neuroprotection studies indicate that NfL has utility as a pharmacodynamic biomarker when measured in the subacute phase after TBI (43, 41), suggesting that NfL should be incorporated as a highly translational pharmacodynamic biomarker when testing new drug treatment strategies for TBI in preclinical models.

SNTF, an N-terminal proteolytic fragment of spectrin, is another emerging blood biomarker of traumatic axonal injury (115). SNTF is generated via calcium overload in injured axons leading to the activation of calpain, which in turn cleaves axonal spectrin (116). Notably, like NfL, SNTF has been shown to preferentially accumulate in axonal swellings in small and large animal models of TBI and in post-mortem examination of human TBI (112,116,117). Surprisingly, however, SNTF and NfL do not co-accumulate in the same axonal swellings, suggesting they may each identify different forms of axonal pathology that evolve at the same time in the white matter (117). While not extensively examined as a blood biomarker in preclinical TBI models, clinical studies of SNTF levels in the blood have shown high sensitivity in predicting which patients with acute concussion will have a poor outcome, such as persisting neurocognitive dysfunction (118–121). Furthermore, blood SNTF levels are associated with white matter abnormalities detected by diffusion MRI, suggesting a loss of brain connectivity via axonal damage (121). Accordingly, blood examination of NfL and SNTF together in preclinical studies may help guide development of therapies specifically targeting traumatic axonal injury (122).

Circulating GFAP levels are sharply increased following moderate-to-severe TBI (47) and smTBI (60,61,79,83,91), with levels doubling from 2 to 4h (47) post-injury (**Fig. 4A,B**). In contrast, there is a delayed peak occurring at 1w following rmTBI (66,92,93) (**Fig. 4C**), which may indicate a dependence of GFAP on the biomechanical load. Most studies report that GFAP blood levels return to sham levels within 10d after TBI, possibly explaining the lack of preclinical TBI studies examining chronic time points. However, in TBI patients, while the initial peak in circulating GFAP overlaps with the pattern observed in rodent TBI models (**Fig. 5**), there is a second peak in humans that occurs many years after TBI (20). Astrocytic endfeet that support BBB integrity can sustain damage following biomechanical impacts, thereby compromising BBB permeability and facilitating the release of GFAP into the circulation (7). Post-traumatic BBB disruption manifests in a biphasic manner, commencing with acute shear stess/injury to endothelial cells in capillaries and arterioles due to the biomechanical damage, followed by a delayed phase characterized by an increase in transcytosis of plasma proteins, contributing to edema (123). Of note, a recent study showed that increased astrocytic reactivity, indicated by elevated plasma GFAP, was associated with proteinopathies involved in AD, suggesting a link between astrocytic reactivity, GFAP levels, and chronic neurodegeneration (124). Additional preclinical studies that include chronic timepoints are needed to determine whether circulating GFAP also displays a biphasic response in rodents, and to understand the source of circulating GFAP. Indeed, preclinical studies present the ideal setting to dissect the relative contribution of GFAP released from perivascular astrocytic endfeet, primarily as a result of a compromised BBB, or from reactive astrocytes.

The prognostic value of GFAP biomarkers has been studied in rodent models, and acute levels of GFAP predict contusion volume (106) and tissue GBDPs (57), a product of activated calpain proteolysis and a selective marker of astrocytic damage in both rodent (125) and human TBI (126). GFAP has been widely used as a pharmacodynamic marker, largely due to the efforts of the Operation Brain Trauma Therapy (OBTT) consortium (127). GFAP levels repeatedly showed a drug treatment effect across injury severities (57,59,65,67,73,74,78,85,93), with only a few exceptions (41,65,82,86). Interestingly, all studies that did not report a reduction in post-traumatic GFAP levels due to drug treatment used either weight-drop (82) or FPI (41,65,86) models to induce TBI. For example, when LEV, the highest scoring drug in the OBTT studies, was administered it produced a reduction in post-traumatic GFAP levels only after CCI, but not in the PBBI (59) nor FPI (41,59) models. Another study reported a trend-towards-reduction in post-traumatic circulating GFAP levels following Glibenclamide administration in the CCI model, but not in the PBBI or FPI models (65). Notably, when LEV was tested in a gyroencephalic FPI model (128) there were no changes in post-traumatic GFAP levels due to LEV treatment. Collectively, these data suggest that post-traumatic GFAP pharmacodynamic response may be model-dependent (59,128).

In preclinical models, t-Tau and p-Tau can be detected in the circulation in the days following TBI. Data in the chronic phase post-injury are scarce, but a progressive increase was observed following moderate-to-severe TBI (**Fig. 4A**) (98,100) and rmTBI (**Fig. 4C**) (46,87,89,93), but not after smTBI (**Fig. 4B**) (55,94,96,101). Notably, one study reported increased t-Tau and p-Tau levels up to one year following rmTBI in tauopathy-prone transgenic TghTau/PS1 mice expressing human tau (97). In TBI patients, t-Tau and p-Tau in blood peaks within 30 days post-injury and subsequently decline towards baseline levels in the subacute phase (7,20,32), before becoming elevated once again during a secondary phase that rebounds and peaks at one year post-injury, possibly due to ongoing white matter injury and chronic neurodegeneration (20,32). Accumulating data suggests that the biomechanical load (either by a single high intensity injury or a milder repeated events) is critical in triggering chronic neurodegenerative processes characterized by Tau pathology, as shown by post-mortem analyses in mice and humans (129,130).

There is little evidence to support the use of circulating tau proteins as prognostic biomarkers for TBI, with only one study showing p-Tau’s prognostic ability to predict post-traumatic seizures in rats (41). t-Tau and p-Tau may hold potential as pharmacodynamic markers, although the evidence is somewhat limited. Some interventions, such as oxygen therapy, have been associated with a temporary reduction in t-Tau protein levels, indicating possible neuroprotective effects (94). Dietary interventions also seem to prevent post-traumatic increases in t-Tau (73,103). Furthermore, a reduction in t-Tau and p-Tau has been reported following treatment with lithium chloride and rosiglitazone in transgenic human tau hTau/PS1 mice subjected to rmTBI (97). However, other treatments such as etanercept and lithium did not alter post-traumatic t-Tau levels (82), nor did treatment with LEV which instead reduced p-Tau levels following TBI (41). While these markers may offer insights into TBI progression and responses to drug treatment, additional studies are clearly needed.

Finally, circulating UCH-L1 levels are modestly impacted by TBI, even after severe TBI in rodents, where UCH-L1 shows a marginal increase early after injury (**Fig. 4A**) (57,59,64,65,69,71,75,85,104,105,131), or no change at all (58,62,70,74,107,108). In addition, the preclinical data indicate that blood UCH-L1 has a weak pharmacodynamic response (65,71,74). In larger animals such as the micro-pig, UCH-L1 levels did not change following mTBI (128). Post-traumatic UCH-L1 responses in rodent TBI models do not mirror the changes observed in human TBI, where UCH-L1 has clinical utility (although it is outperformed by GFAP in mTBI (132)). It will be important to determine whether this discrepancy is attributed to species-specific variations in protein biomarker levels or involves other mechanisms.

## Can biomarker trajectories provide anchor points that will enhance translational research?

The reliability of preclinical models hinges on their ability to replicate the evolution of structural and functional abnormalities associated with TBI in patients. Thus, rodent models become essential for studying the pathobiology underpinning those changes and for exploring therapeutic interventions. However, pathophysiological processes, such as axonal damage and neuroinflammation, operate on distinct timescales in rodents and humans, and there is no universal ‘conversion rate’ between these processes (111). Incorporating longitudinal assessments of TBI biomarkers in preclinical research may provide valuable insights into these differences. In **Fig. 5** we attempt to align the temporal dynamics of the most well-characterized biomarkers in the preclinical setting, NfL and GFAP between rodents and humans. During the acute phase following injury, there is a similarity in the time-to-first peak for GFAP in rodents (24 hours) and humans (24 hours). For NfL, however, the initial increase occurs at 3 days post-injury in rodents and at 14-21 days post-injury in humans. Notably, data available for the chronic phase suggest the conversion rate for NfL remains stable over time with the time-to-decrease nearly six times slower in humans (6w) compared to rodents (1w). The fact that conversion rates may be unique to each biomarker suggests they could also help in the choice of appropriate models based on targeted mechanisms.

From a methodological perspective, the use of comparable analysis platforms is key to limiting variability between preclinical and clinical data. The surge in clinical protein TBI biomarker data has been made possible in large part through the development of sensitive immunoassays that have a high degree of analytical validation, including measures of precision, robustness, dilution linearity, upper and lower limits of detection, and coefficients of variation. It is important to mention that assays used clinically may not always be translatable to rodent models, particularly if the primary antibody used in the clinical assay does not cross-react with its murine counterpart (*e.g.,* GFAP, Tau). Therefore, there is need to develop additional biomarker assays for use in rodents that have rigorous analytical validation, low volume requirements, and sensitivity that parallels the established clinical assays.

To address these variables and foster greater alignment, collaborative efforts to share preclinical and clinical data and biospecimens are needed. In this context the establishment of a virtual preclinical biobank within new or existing preclinical and clinical consortia (i.e. the International Initiative for Traumatic Brain Injury Research, InTBIR) would foster centralized, rigorous, analyses of preclinical TBI biomarkers. It is anticipated that such a research platform would provide the missing anchor points between preclinical and clinical studies, which will greatly enhance the overall quality and translational potential of research in the TBI field.

## Knowledge gaps, limitations and conclusions

This review highlights the need for additional preclinical data on the subacute and chronic phases of TBI that are critical to fully understand the biomarkers’ translational potential. Experiments exploring the effect of sex (133) and age on the levels and kinetics of blood biomarkers following TBI are currently lacking. The majority of studies have evaluated preclinical TBI biomarkers exclusively in male mice; a disparity mirrored in large clinical studies (*e.g.,* 6,105). Similarly, few studies have examined circulating biomarker changes in geriatric or pediatric TBI models. This is particularly noteworthy considering that the elderly population registers the highest frequency of TBI-related hospital admissions (135) and have significantly poorer outcomes compared to young patients with similar injury severity (136). This pattern is mirrored in aged TBI mice (137). Notably, blood NfL levels exhibit an age-related increase in TBI patients, hinting at age-linked vulnerability to neuronal damage (136). Determining whether this susceptibility is attributed to age itself or to age-related comorbidities presents challenges in the clinical context. Animal models of TBI offer an opportunity to disentangle these complex mechanisms and shed light on their interactions. Additionally, in the pediatric population, there is a significant value in avoiding CT/MRI scans through the use of biomarkers; further studies in juvenile rodent models will be required to fill in this major research gap. Addressing these knowledge gaps holds the key to advancing our comprehension of TBI pathophysiology and, ultimately, improving patient outcomes.

This systematic review is not without limitations. Firstly, while we focused on the most promising biomarkers of TBI from a translational point of view, the panel of biomarkers investigated is not exhaustive. Other biomarker types, such as miRNAs and different proteins, merit similar attention in future research endeavors. Secondly, our data analysis involved categorizing studies based on injury severity, which ranged from ‘mild’ to ‘moderate-to-severe’, as defined by the authors. It is important to acknowledge that this terminology holds subjectivity. Indeed, efforts are underway to establish a more standardized classification system for TBI that offers an objective representation of injury severity and disease trajectories. In the context of preclinical studies, maintaining alignment with clinical definitions is crucial, and the utilization of blood biomarkers could prove instrumental in facilitating translation within this domain.

In conclusion, preclinical models replicate the pattern and trajectories of blood biomarkers that are observed in human TBI. Preclinical biomarkers also have the potential to bridge the gap between preclinical and clinical research that is needed to develop new therapeutic interventions for TBI. This is particularly noteworthy for biomarkers such as NfL, which demonstrates predictive value for long-term clinically relevant outcomes. Moreover, NfL along with GFAP hold pharmacodynamic potential, showing changes in response to therapeutic interventions. By including longitudinal assessment of circulating biomarkers in preclinical drug screening the development pipeline will be improved, which will increase the likelihood of translating preclinical discoveries into new therapeutics with clinical efficacy.

## Abbreviations

NfL: neurofilament-light
t-Tau: total Tau
p-Tau: phosphorylated-Tau
UCH-L1: ubiquitin C-terminal hydrolase
GFAP: glial fibrillary acidic protein
TBI: traumatic brain injury
smTBI: single mild TBI
rmTBI: repetitive mild TBI
h: hours
d: days
w: weeks
m: months

## References

1. Maas AIR, Menon DK, Manley GT, Abrams M, Åkerlund C, Andelic N, et al. Traumatic brain injury: progress and challenges in prevention, clinical care, and research. Lancet Neurol. 2022 Nov;21(11):1004–60.

2. Majdan M, Plancikova D, Brazinova A, Rusnak M, Nieboer D, Feigin V, et al. Epidemiology of traumatic brain injuries in Europe: a cross-sectional analysis. Lancet Public Health. 2016 Dec;1(2):e76–83.

3. Wang KK, Yang Z, Zhu T, Shi Y, Rubenstein R, Tyndall JA, et al. An update on diagnostic and prognostic biomarkers for traumatic brain injury. Expert Rev Mol Diagn. 2018 Feb;18(2):165–80.

4. Åkerlund CAI, Holst A, Stocchetti N, Steyerberg EW, Menon DK, Ercole A, et al. Clustering identifies endotypes of traumatic brain injury in an intensive care cohort: a CENTER-TBI study. Crit Care. 2022 Dec;26(1):228.

5. Czeiter E, Amrein K, Gravesteijn BY, Lecky F, Menon DK, Mondello S, et al. Blood biomarkers on admission in acute traumatic brain injury: Relations to severity, CT findings and care path in the CENTER-TBI study. EBioMedicine. 2020 Jun;56:102785.

6. Newcombe VFJ, Ashton NJ, Posti JP, Glocker B, Manktelow A, Chatfield DA, et al. Post-acute blood biomarkers and disease progression in traumatic brain injury. Brain. 2022 Jun;145(6):2064–76.

7. Graham NSN, Zimmerman KA, Moro F, Heslegrave A, Maillard SA, Bernini A, et al. Axonal marker neurofilament light predicts long-term outcomes and progressive neurodegeneration after traumatic brain injury. Sci Transl Med. 2021 Sep;13(613):eabg9922.

8. Papa L, Ladde JG, O’Brien JF, Thundiyil JG, Tesar J, Leech S, et al. Evaluation of Glial and Neuronal Blood Biomarkers Compared With Clinical Decision Rules in Assessing the Need for Computed Tomography in Patients With Mild Traumatic Brain Injury. JAMA Netw Open. 2022 Mar;5(3):e221302.

9. Yue JK, Yuh EL, Korley FK, Winkler EA, Sun X, Puffer RC, et al. Association between plasma GFAP concentrations and MRI abnormalities in patients with CT-negative traumatic brain injury in the TRACK-TBI cohort: a prospective multicentre study. Lancet Neurol. 2019 Oct;18(10):953–61.

10. Helmrich IRAR, Czeiter E, Amrein K, Büki A, Lingsma HF, Menon DK, et al. Incremental prognostic value of acute serum biomarkers for functional outcome after traumatic brain injury (CENTER-TBI): an observational cohort study. Lancet Neurol. 2022 Sep;21(9):792–802.

11. Korley FK, Jain S, Sun X, Puccio AM, Yue JK, Gardner RC, et al. Prognostic value of day-of-injury plasma GFAP and UCH-L1 concentrations for predicting functional recovery after traumatic brain injury in patients from the US TRACK-TBI cohort: an observational cohort study. Lancet Neurol. 2022 Sep;21(9):803–13.

12. Scott G, Zetterberg H, Jolly A, Cole JH, De Simoni S, Jenkins PO, et al. Minocycline reduces chronic microglial activation after brain trauma but increases neurodegeneration. Brain. 2018 Feb;141(2):459–71.

13. Zanier ER, Pischiutta F, Rulli E, Vargiolu A, Elli F, Gritti P, et al. MesenchymAl stromal cells for Traumatic bRain Injury (MATRIx): a study protocol for a multicenter, double-blind, randomised, placebo-controlled phase II trial. Intensive Care Med Exp. 2023 Aug;11(1):56.

14. Zetterberg H, Blennow K. Fluid biomarkers for mild traumatic brain injury and related conditions. Nat Rev Neurol. 2016 Oct;12(10):563–74.

15. Gaetani L, Blennow K, Calabresi P, Di Filippo M, Parnetti L, Zetterberg H. Neurofilament light chain as a biomarker in neurological disorders. J Neurol Neurosurg Psychiatry. 2019 Aug;90(8):870–81.

16. Donat CK, Yanez Lopez M, Sastre M, Baxan N, Goldfinger M, Seeamber R, et al. From biomechanics to pathology: predicting axonal injury from patterns of strain after traumatic brain injury. Brain. 2021 Feb;144(1):70–91.

17. Uryu K, Chen XH, Martinez D, Browne KD, Johnson VE, Graham DI, et al. Multiple proteins implicated in neurodegenerative diseases accumulate in axons after brain trauma in humans. Exp Neurol. 2007 Dec;208(2):185–92.

18. Bacioglu M, Maia LF, Preische O, Schelle J, Apel A, Kaeser SA, et al. Neurofilament Light Chain in Blood and CSF as Marker of Disease Progression in Mouse Models and in Neurodegenerative Diseases. Neuron. 2016 Jul;91(2):494–6.

19. Bäckström D, Linder J, Jakobson Mo S, Riklund K, Zetterberg H, Blennow K, et al. NfL as a biomarker for neurodegeneration and survival in Parkinson disease. Neurology. 2020 Aug;95(7):e827–38.

20. Shahim P, Politis A, van der Merwe A, Moore B, Ekanayake V, Lippa SM, et al. Time course and diagnostic utility of NfL, tau, GFAP, and UCH-L1 in subacute and chronic TBI. Neurology. 2020 Aug;95(6):e623–36.

21. Eng LF, Ghirnikar RS, Lee YL. Glial fibrillary acidic protein: GFAP-thirty-one years (1969-2000). Neurochem Res. 2000 Oct;25(9/10):1439–51.

22. Yang Z, Wang KKW. Glial fibrillary acidic protein: from intermediate filament assembly and gliosis to neurobiomarker. Trends Neurosci. 2015 Jun;38(6):364–74.

23. Janigro D, Mondello S, Posti JP, Unden J. GFAP and S100B: What You Always Wanted to Know and Never Dared to Ask. Front Neurol. 2022 Mar;13:835597.

24. Lei J, Gao G, Feng J, Jin Y, Wang C, Mao Q, et al. Glial fibrillary acidic protein as a biomarker in severe traumatic brain injury patients: a prospective cohort study. Crit Care. 2015 Dec;19(1):362.

25. Vos PE, Lamers KJB, Hendriks JCM, van Haaren M, Beems T, Zimmerman C, et al. Glial and neuronal proteins in serum predict outcome after severe traumatic brain injury. Neurology. 2004 Apr;62(8):1303–10.

26. Okonkwo DO, Puffer RC, Puccio AM, Yuh EL, Yue JK, Diaz-Arrastia R, et al. Point-of-Care Platform Blood Biomarker Testing of Glial Fibrillary Acidic Protein versus S100 Calcium-Binding Protein B for Prediction of Traumatic Brain Injuries: A Transforming Research and Clinical Knowledge in Traumatic Brain Injury Study. J Neurotrauma. 2020 Dec;37(23):2460–7.

27. Papa L, Silvestri S, Brophy GM, Giordano P, Falk JL, Braga CF, et al. GFAP Out-Performs S100β in Detecting Traumatic Intracranial Lesions on Computed Tomography in Trauma Patients with Mild Traumatic Brain Injury and Those with Extracranial Lesions. J Neurotrauma. 2014 Nov;31(22):1815–22.

28. Barbier P, Zejneli O, Martinho M, Lasorsa A, Belle V, Smet-Nocca C, et al. Role of Tau as a Microtubule-Associated Protein: Structural and Functional Aspects. Front Aging Neurosci. 2019 Aug;11:204.

29. McKee AC, Stern RA, Nowinski CJ, Stein TD, Alvarez VE, Daneshvar DH, et al. The spectrum of disease in chronic traumatic encephalopathy. Brain. 2013 Jan;136(Pt 1):43–64.

30. Turk KW, Geada A, Alvarez VE, Xia W, Cherry JD, Nicks R, et al. A comparison between tau and amyloid-β cerebrospinal fluid biomarkers in chronic traumatic encephalopathy and Alzheimer disease. Alzheimers Res Ther. 2022 Feb;14(1):28.

31. Milà-Alomà M, Ashton NJ, Shekari M, Salvadó G, Ortiz-Romero P, Montoliu-Gaya L, et al. Plasma p-tau231 and p-tau217 as state markers of amyloid-β pathology in preclinical Alzheimer’s disease. Nat Med. 2022 Sep;28(9):1797–1801.

32. Rubenstein R, Chang B, Yue JK, Chiu A, Winkler EA, Puccio AM, et al. Comparing Plasma Phospho Tau, Total Tau, and Phospho Tau-Total Tau Ratio as Acute and Chronic Traumatic Brain Injury Biomarkers. JAMA Neurol. 2017 Sep;74(9):1063–72.

33. Rubenstein R, McQuillan L, Wang KKW, Robertson C, Chang B, Yang Z, et al. Temporal Profiles of P-Tau, T-Tau, and P-Tau:Tau Ratios in Cerebrospinal Fluid and Blood from Moderate-Severe Traumatic Brain Injury Patients and Relationship to 6-12 Month Global Outcomes. J Neurotrauma. 2023 Nov; Epub ahead of print.

34. Gonzalez-Ortiz F, Dulewicz M, Ashton NJ, Kac PR, Zetterberg H, Andersson E, et al. Association of Serum Brain-Derived Tau With Clinical Outcome and Longitudinal Change in Patients With Severe Traumatic Brain Injury. JAMA Netw Open. 2023 Jul;6(7):e2321554.

35. Wang KKW, Kobeissy FH, Shakkour Z, Tyndall JA. Thorough overview of ubiquitin C-terminal hydrolase-L1 and glial fibrillary acidic protein as tandem biomarkers recently cleared by US Food and Drug Administration for the evaluation of intracranial injuries among patients with traumatic brain injury. Acute Med Surg. 2021;8(1):e622.

36. Sokka AL, Putkonen N, Mudo G, Pryazhnikov E, Reijonen S, Khiroug L, et al. Endoplasmic Reticulum Stress Inhibition Protects against Excitotoxic Neuronal Injury in the Rat Brain. J Neurosci. 2007 Jan;27(4):901–8.

37. Papa L, Lewis LM, Silvestri S, Falk JL, Giordano P, Brophy GM, et al. Serum levels of ubiquitin C-terminal hydrolase distinguish mild traumatic brain injury from trauma controls and are elevated in mild and moderate traumatic brain injury patients with intracranial lesions and neurosurgical intervention. J Trauma Acute Care Surg. 2012 May;72(5):1335–44.

38. Macleod MR, O’Collins T, Howells DW, Donnan GA. Pooling of Animal Experimental Data Reveals Influence of Study Design and Publication Bias. Stroke. 2004 May;35(5):1203–8.

39. Li X, Pierre K, Yang Z, Nguyen L, Johnson G, Venetucci J, et al. Blood-Based Brain and Global Biomarker Changes after Combined Hypoxemia and Hemorrhagic Shock in a Rat Model of Penetrating Ballistic-Like Brain Injury. Neurotrauma Rep. 2021 Aug;2(1):370–80.

40. Heiskanen M, Jääskeläinen O, Manninen E, Das Gupta S, Andrade P, Ciszek R, et al. Plasma Neurofilament Light Chain (NF-L) Is a Prognostic Biomarker for Cortical Damage Evolution but Not for Cognitive Impairment or Epileptogenesis Following Experimental TBI. Int J Mol Sci. 2022 Dec;23(23):15208.

41. Saletti PG, Mowrey WB, Liu W, Li Q, McCullough J, Aniceto R, et al. Early preclinical plasma protein biomarkers of brain trauma are influenced by early seizures and levetiracetam. Epilepsia Open. 2023 Jun;8(2):586–608.

42. Boucher ML, Conley G, Nowlin J, Qiu J, Kawata K, Bazarian JJ, et al. Titrating the Translational Relevance of a Low-Level Repetitive Head Impact Model. Front Neurol. 2022 Jun;13:857654.

43. Cheng WH, Stukas S, Martens KM, Namjoshi DR, Button EB, Wilkinson A, et al. Age at injury and genotype modify acute inflammatory and neurofilament-light responses to mild CHIMERA traumatic brain injury in wild-type and APP/PS1 mice. Exp Neurol. 2018 Mar;301:26–38.

44. Dickstein DL, De Gasperi R, Gama Sosa MA, Perez-Garcia G, Short JA, Sosa H, et al. Brain and blood biomarkers of tauopathy and neuronal injury in humans and rats with neurobehavioral syndromes following blast exposure. Mol Psychiatry. 2021 Oct;26(10):5940–54.

45. Moro F, Lisi I, Tolomeo D, Vegliante G, Pascente R, Mazzone E, et al. Acute Blood Levels of Neurofilament Light Indicate One-Year White Matter Pathology and Functional Impairment in Repetitive Mild Traumatic Brain Injured Mice. J Neurotrauma. 2023 Jun;40(11–12):1144–63.

46. O’Brien WT, Pham L, Brady RD, Bain J, Yamakawa GR, Sun M, et al. Temporal profile and utility of serum neurofilament light in a rat model of mild traumatic brain injury. Exp Neurol. 2021 Jul;341:113698.

47. O’Brien WT, Wright DK, van Emmerik ALJJ, Bain J, Brkljaca R, Christensen J, et al. Serum neurofilament light as a biomarker of vulnerability to a second mild traumatic brain injury. Transl Res. 2023 May;255:77–84.

48. Pham L, Wright DK, O’Brien WT, Bain J, Huang C, Sun M, et al. Behavioral, axonal, and proteomic alterations following repeated mild traumatic brain injury: Novel insights using a clinically relevant rat model. Neurobiol Dis. 2021 Jan;148:105151.

49. Stemper BD, Shah A, Chiariello R, McCarthy C, Jessen K, Sarka B, et al. A Preclinical Rodent Model for Repetitive Subconcussive Head Impact Exposure in Contact Sport Athletes. Front Behav Neurosci. 2022 Mar;16:805124.

50. Bashir A, Abebe ZA, McInnes KA, Button EB, Tatarnikov I, Cheng WH, et al. Increased severity of the CHIMERA model induces acute vascular injury, sub-acute deficits in memory recall, and chronic white matter gliosis. Exp Neurol. 2020 Feb;324:113116.

51. Diomede L, Zanier ER, Moro F, Vegliante G, Colombo L, Russo L, et al. Aβ1-6A2V(D) peptide, effective on Aβ aggregation, inhibits tau misfolding and protects the brain after traumatic brain injury. Mol Psychiatry. 2023 Jun;28(6):2433–2444.

52. Wong KR, O’Brien WT, Sun M, Yamakawa G, O’Brien TJ, Mychasiuk R, et al. Serum Neurofilament Light as a Biomarker of Traumatic Brain Injury in the Presence of Concomitant Peripheral Injury. Biomark Insights. 2021 Oct;16:11772719211053448.

53. Thau-Zuchman O, Ingram R, Harvey GG, Cooke T, Palmas F, Pallier PN, et al. A Single Injection of Docosahexaenoic Acid Induces a Pro-Resolving Lipid Mediator Profile in the Injured Tissue and a Long-Lasting Reduction in Neurological Deficit after Traumatic Brain Injury in Mice. J Neurotrauma. 2020 Jan;37(1):66–79.

54. Koza LA, Pena C, Russell M, Smith AC, Molnar J, Devine M, et al. Immunocal® limits gliosis in mouse models of repetitive mild-moderate traumatic brain injury. Brain Res. 2023 Jun;1808:148338.

55. Ahmed F, Plantman S, Cernak I, Agoston DV. The Temporal Pattern of Changes in Serum Biomarker Levels Reveals Complex and Dynamically Changing Pathologies after Exposure to a Single Low-Intensity Blast in Mice. Front Neurol. 2015 Jun;6:114.

56. Arun P, Abu-Taleb R, Oguntayo S, Tanaka M, Wang Y, Valiyaveettil M, et al. Distinct patterns of expression of traumatic brain injury biomarkers after blast exposure: Role of compromised cell membrane integrity. Neurosci Lett. 2013 Sep;552:87–91.

57. Boutté AM, Deng-Bryant Y, Johnson D, Tortella FC, Dave JR, Shear DA, et al. Serum Glial Fibrillary Acidic Protein Predicts Tissue Glial Fibrillary Acidic Protein Break-Down Products and Therapeutic Efficacy after Penetrating Ballistic-Like Brain Injury. J Neurotrauma. 2016 Jan;33(1):147–56.

58. Bramlett HM, Dietrich WD, Dixon CE, Shear DA, Schmid KE, Mondello S, et al. Erythropoietin Treatment in Traumatic Brain Injury: Operation Brain Trauma Therapy. J Neurotrauma. 2016 Mar;33(6):538–52.

59. Browning M, Shear DA, Bramlett HM, Dixon CE, Mondello S, Schmid KE, et al. Levetiracetam Treatment in Traumatic Brain Injury: Operation Brain Trauma Therapy. J Neurotrauma. 2016 Mar;33(6):581–94.

60. Candy S, Ma I, McMahon JM, Farrell M, Mychasiuk R. Staying in the game: a pilot study examining the efficacy of protective headgear in an animal model of mild traumatic brain injury (mTBI). Brain Inj. 2017 Sep;31(11):1521–9.

61. Cikriklar HI, Ekici MA, Ozbek Z, Cosan DT, Baydemir C, Yürümez Y. Investigation of the course of GFAP in blood in the initial 24 hours in rats subjected to minor head trauma. Eur Rev Med Pharmacol Sci. 2013 Dec;17(24):3391–7.

62. Dixon CE, Bramlett HM, Dietrich WD, Shear DA, Yan HQ, Deng-Bryant Y, et al. Cyclosporine Treatment in Traumatic Brain Injury: Operation Brain Trauma Therapy. J Neurotrauma. 2016 Mar;33(6):553–66.

63. Hiskens MI, Schneiders AG, Vella RK, Fenning AS. Repetitive mild traumatic brain injury affects inflammation and excitotoxic mRNA expression at acute and chronic time-points. PLoS One. 2021 May;16(5):e0251315.

64. Huang X jian, Glushakova O, Mondello S, Van K, Hayes RL, Lyeth BG. Acute Temporal Profiles of Serum Levels of UCH-L1 and GFAP and Relationships to Neuronal and Astroglial Pathology following Traumatic Brain Injury in Rats. J Neurotrauma. 2015 Aug;32(16):1179–89.

65. Jha RM, Mondello S, Bramlett HM, Dixon CE, Shear DA, Dietrich WD, et al. Glibenclamide Treatment in Traumatic Brain Injury: Operation Brain Trauma Therapy. J Neurotrauma. 2021 Mar;38(5):628–45.

66. Kamnaksh A, Kwon SK, Kovesdi E, Ahmed F, Barry ES, Grunberg NE, et al. Neurobehavioral, cellular, and molecular consequences of single and multiple mild blast exposure. Electrophoresis. 2012 Dec;33(24):3680–92.

67. Kim CK, Park JS, Kim E, Oh MK, Lee YT, Yoon KJ, et al. The effects of early exercise in traumatic brain-injured rats with changes in motor ability, brain tissue, and biomarkers. BMB Rep. 2022 Oct;55(10):512–7.

68. Liu MD, Luo P, Wang ZJ, Fei Z. Changes of serum Tau, GFAP, TNF-α and malonaldehyde after blast-related traumatic brain injury. Chin J Traumatol Zhonghua Chuang Shang Za Zhi. 2014;17(6):317–22.

69. Mondello S, Shear DA, Bramlett HM, Dixon CE, Schmid KE, Dietrich WD, et al. Insight into Pre-Clinical Models of Traumatic Brain Injury Using Circulating Brain Damage Biomarkers: Operation Brain Trauma Therapy. J Neurotrauma. 2016 Mar;33(6):595–605.

70. Mountney A, Bramlett HM, Dixon CE, Mondello S, Dietrich WD, Wang KKW, et al. Simvastatin Treatment in Traumatic Brain Injury: Operation Brain Trauma Therapy. J Neurotrauma. 2016 Mar;33(6):567–80.

71. Osier ND, Bramlett HM, Shear DA, Mondello S, Carlson SW, Dietrich WD, et al. Kollidon VA64 Treatment in Traumatic Brain Injury: Operation Brain Trauma Therapy. J Neurotrauma. 2021 Sep;38(17):2454–72.

72. Rubenstein R, Chang B, Grinkina N, Drummond E, Davies P, Ruditzky M, et al. Tau phosphorylation induced by severe closed head traumatic brain injury is linked to the cellular prion protein. Acta Neuropathol Commun. 2017 Dec;5(1):30.

73. Scrimgeour AG, Condlin ML, Loban A, DeMar JC. Omega-3 Fatty Acids and Vitamin D Decrease Plasma T-Tau, GFAP, and UCH-L1 in Experimental Traumatic Brain Injury. Front Nutr. 2021 Jun;8:685220.

74. Shear DA, Dixon CE, Bramlett HM, Mondello S, Dietrich WD, Deng-Bryant Y, et al. Nicotinamide Treatment in Traumatic Brain Injury: Operation Brain Trauma Therapy. J Neurotrauma. 2016 Mar;33(6):523–37.

75. Svetlov SI, Prima V, Kirk DR, Gutierrez H, Curley KC, Hayes RL, et al. Morphologic and Biochemical Characterization of Brain Injury in a Model of Controlled Blast Overpressure Exposure. J Trauma Inj Infect Crit Care. 2010 Oct;69(4):795–804.

76. Svetlov SI. Neuro-glial and systemic mechanisms of pathological responses in rat models of primary blast overpressure compared to “composite” blast. Front Neurol. 2012 Feb;3:15.

77. VandeVord PJ, Sajja VSSS, Ereifej E, Hermundstad A, Mao S, Hadden TJ. Chronic Hormonal Imbalance and Adipose Redistribution Is Associated with Hypothalamic Neuropathology following Blast Exposure. J Neurotrauma. 2016 Jan;33(1):82–8.

78. Wei C, Luo Y, Peng L, Huang Z, Pan Y. Expression of Notch and Wnt/β-catenin signaling pathway in acute phase severe brain injury rats and the effect of exogenous thyroxine on those pathways. Eur J Trauma Emerg Surg. 2021 Dec;47(6):2001–2015.

79. Yang SH, Gustafson J, Gangidine M, Stepien D, Schuster R, Pritts TA, et al. A murine model of mild traumatic brain injury exhibiting cognitive and motor deficits. J Surg Res. 2013 Oct;184(2):981–8.

80. Button EB, Cheng WH, Barron C, Cheung H, Bashir A, Cooper J, et al. Development of a novel, sensitive translational immunoassay to detect plasma glial fibrillary acidic protein (GFAP) after murine traumatic brain injury. Alzheimers Res Ther. 2021 Dec;13(1):58.

81. Cikriklar HI, Uysal O, Ekici MA, Ozbek Z, Cosan DT, Yucel M, et al. Effectiveness of GFAP in Determining Neuronal Damage in Rats with Induced Head Trauma. Turk Neurosurg. 2016;26(6):878–89.

82. Ekici MA, Uysal O, Cikriklar HI, Özbek Z, Turgut Cosan D, Baydemir C, et al. Effect of etanercept and lithium chloride on preventing secondary tissue damage in rats with experimental diffuse severe brain injury. Eur Rev Med Pharmacol Sci. 2014 Oct;18(1):10–27.

83. Li Y, Zhang L, Kallakuri S, Cohen A, Cavanaugh JM. Correlation of mechanical impact responses and biomarker levels: A new model for biomarker evaluation in TBI. J Neurol Sci. 2015 Dec;359(1–2):280–6.

84. Gao N, Zhang-Brotzge X, Wali B, Sayeed I, Chern JJ, Blackwell LS, et al. Plasma osteopontin may predict neuroinflammation and the severity of pediatric traumatic brain injury. J Cereb Blood Flow Metab. 2020 Jan;40(1):35–43.

85. Kobeissy F, Mallah K, Zibara K, Dakroub F, Dalloul Z, Nasser M, et al. The effect of clopidogrel and aspirin on the severity of traumatic brain injury in a rat model. Neurochem Int. 2022 Mar;154:105301.

86. Liang DY, Sahbaie P, Sun Y, Irvine KA, Shi X, Meidahl A, et al. TBI-induced nociceptive sensitization is regulated by histone acetylation. IBRO Rep. 2017 Jun;2:14–23.

87. Robinson S, Winer JL, Berkner J, Chan LAS, Denson JL, Maxwell JR, et al. Imaging and serum biomarkers reflecting the functional efficacy of extended erythropoietin treatment in rats following infantile traumatic brain injury. J Neurosurg Pediatr. 2016 Jun;17(6):739–55.

88. Tadepalli SA, Bali ZK, Bruszt N, Nagy LV, Amrein K, Fazekas B, et al. Long-term cognitive impairment without diffuse axonal injury following repetitive mild traumatic brain injury in rats. Behav Brain Res. 2020 Jan;378:112268.

89. Pierce JD, Gupte R, Thimmesch A, Shen Q, Hiebert JB, Brooks WM, et al. Ubiquinol treatment for TBI in male rats: Effects on mitochondrial integrity, injury severity, and neurometabolism. J Neurosci Res. 2018 Jun;96(6):1080–92.

90. Kmeťová K, Drobná D, Lipták R, Hodosy J, Celec P. Early dynamics of glial fibrillary acidic protein and extracellular DNA in plasma of mice after closed head traumatic brain injury. Neurochirurgie. 2022 Dec;68(6):e68–74.

91. Plog BA, Dashnaw ML, Hitomi E, Peng W, Liao Y, Lou N, et al. Biomarkers of Traumatic Injury Are Transported from Brain to Blood via the Glymphatic System. J Neurosci. 2015 Jan;35(2):518–26.

92. Ahmed FA, Kamnaksh A, Kovesdi E, Long JB, Agoston DV. Long-term consequences of single and multiple mild blast exposure on select physiological parameters and blood-based biomarkers. Electrophoresis. 2013 Aug;34(15):2229–33.

93. Pozdnyakov DI, Miroshnichenko KA, Voronkov AV, Kovaleva TG. The Administration of the New Pyrimidine Derivative-4-{2-[2-(3,4-Dimethoxyphenyl)-Vinyl]-6-Ethyl-4-Oxo-5-Phenyl-4H-Pyrimidine-1-Il}Benzsulfamide Restores the Activity of Brain Cells in Experimental Chronic Traumatic Encephalopathy by Maintaining Mitochondrial Function. Medicina (Kaunas). 2019 Jul;55(7):386.

94. Li Y, Lv W, Cheng G, Wang S, Liu B, Zhao H, et al. Effect of Early Normobaric Hyperoxia on Blast-Induced Traumatic Brain Injury in Rats. Neurochem Res. 2020 Nov;45(11):2723–31.

95. Liliang PC, Liang CL, Lu K, Wang KW, Weng HC, Hsieh CH, et al. Relationship between injury severity and serum tau protein levels in traumatic brain injured rats. Resuscitation. 2010 Sep;81(9):1205–8.

96. Rostami E, Davidsson J, Ng KC, Lu J, Gyorgy A, Walker J, et al. A Model for Mild Traumatic Brain Injury that Induces Limited Transient Memory Impairment and Increased Levels of Axon Related Serum Biomarkers. Front Neurol. 2012 Jul;3:115.

97. Rubenstein R, Sharma DR, Chang B, Oumata N, Cam M, Vaucelle L, et al. Novel Mouse Tauopathy Model for Repetitive Mild Traumatic Brain Injury: Evaluation of Long-Term Effects on Cognition and Biomarker Levels After Therapeutic Inhibition of Tau Phosphorylation. Front Neurol. 2019 Mar;10:124.

98. Rubenstein R, Chang B, Davies P, Wagner AK, Robertson CS, Wang KKW. A Novel, Ultrasensitive Assay for Tau: Potential for Assessing Traumatic Brain Injury in Tissues and Biofluids. J Neurotrauma. 2015 Mar;32(5):342–52.

99. Yang Z, Wang P, Morgan D, Lin D, Pan J, Lin F, et al. Temporal MRI characterization, neurobiochemical and neurobehavioral changes in a mouse repetitive concussive head injury model. Sci Rep. 2015 Jun;5:11178.

100. Cheng WH, Cheung H, Kang A, Fan J, Cooper J, Anwer M, et al. Altered Tau Kinase Activity in rTg4510 Mice after a Single Interfaced CHIMERA Traumatic Brain Injury. Int J Mol Sci. 2023 May;24(11):9439.

101. Tomita K, Nakada Taki, Oshima T, Kawaguchi R, Oda S. Serum levels of tau protein increase according to the severity of the injury in DAI rat model. F1000Research. 2020 Jan;9:29.

102. Yao C, Williams AJ, Ottens AK, May Lu XC, Chen R, Wang KK, et al. Detection of protein biomarkers using high-throughput immunoblotting following focal ischemic or penetrating ballistic-like brain injuries in rats. Brain Inj. 2008 Sep;22(10):723–32.

103. Siahaan AMP, Japardi I, Rambe AS, Indharty RS, Ichwan M. Turmeric Extract Supplementation Reduces Tau Protein Level in Repetitive Traumatic Brain Injury Model. Open Access Maced J Med Sci. 2018 Nov;6(11):1953–8.

104. Aleem M, Goswami N, Manda K. Serum biomarkers based neurotrauma severity scale: a study in the mice model of fluid percussion injury. Acta Neurobiol Exp. 2022 Jul;82(2):147–156.

105. Liu MC, Akinyi L, Scharf D, Mo J, Larner SF, Muller U, et al. Ubiquitin C-terminal hydrolase-L1 as a biomarker for ischemic and traumatic brain injury in rats. Eur J Neurosci. 2010 Feb;31(4):722–32.

106. Mondello S, Shear DA, Bramlett HM, Dixon CE, Schmid KE, Dietrich WD, et al. Insight into Pre-Clinical Models of Traumatic Brain Injury Using Circulating Brain Damage Biomarkers: Operation Brain Trauma Therapy. J Neurotrauma. 2016 Mar;33(6):595–605.

107. Wallen TE, Singer KE, Elson NC, Baucom MR, England LG, Schuster RM, et al. Defining Endotheliopathy in Murine Polytrauma Models. Shock. 2022 Jun;57(6):291–8.

108. Morris MC, Bercz A, Niziolek GM, Kassam F, Veile R, Friend LA, et al. UCH-L1 is a Poor Serum Biomarker of Murine Traumatic Brain Injury After Polytrauma. J Surg Res. 2019 Dec;244:63–8.

109. Shin SS, Hefti MM, Mazandi VM, Issadore DA, Meaney DF, Schneider ALC, et al. Plasma Neurofilament Light and Glial Fibrillary Acidic Protein Levels over Thirty Days in a Porcine Model of Traumatic Brain Injury. J Neurotrauma. 2022 Jul;39(13–14):935–43.

110. Millecamps S, Gowing G, Corti O, Mallet J, Julien JP. Conditional NF-L Transgene Expression in Mice for *In Vivo* Analysis of Turnover and Transport Rate of Neurofilaments. J Neurosci. 2007 May;27(18):4947–56.

111. Agoston DV, Vink R, Helmy A, Risling M, Nelson D, Prins M. How to Translate Time: The Temporal Aspects of Rodent and Human Pathobiological Processes in Traumatic Brain Injury. J Neurotrauma. 2019 Jun;36(11):1724–37.

112. Johnson VE, Weber MT, Xiao R, Cullen DK, Meaney DF, Stewart W, et al. Mechanical disruption of the blood-brain barrier following experimental concussion. Acta Neuropathol. 2018 May;135(5):711–26.

113. Sandelius Å, Zetterberg H, Blennow K, Adiutori R, Malaspina A, Laura M, et al. Plasma neurofilament light chain concentration in the inherited peripheral neuropathies. Neurology. 2018 Feb;90(6):e518–24.

114. Ladang A, Kovacs S, Lengelé L, Locquet M, Reginster JY, Bruyère O, et al. Neurofilament light chain concentration in an aging population. Aging Clin Exp Res. 2022 Feb;34(2):331–9.

115. Zetterberg H, Smith DH, Blennow K. Biomarkers of mild traumatic brain injury in cerebrospinal fluid and blood. Nat Rev Neurol. 2013 Apr;9(4):201–10.

116. Büki A, Siman R, Trojanowski JQ, Povlishock JT. The Role of Calpail-Mediated Spectrin Proteolysis in Traumatically Induced Axonal Injury: J Neuropathol Exp Neurol. 1999 Apr;58(4):365–75.

117. Saatman KE, Abai B, Grosvenor A, Vorwerk CK, Smith DH, Meaney DF. Traumatic Axonal Injury Results in Biphasic Calpain Activation and Retrograde Transport Impairment in Mice. J Cereb Blood Flow Metab. 2003 Jan;23(1):34–42.

118. Siman R, Shahim P, Tegner Y, Blennow K, Zetterberg H, Smith DH. Serum SNTF Increases in Concussed Professional Ice Hockey Players and Relates to the Severity of Postconcussion Symptoms. J Neurotrauma. 2015 Sep;32(17):1294–300.

119. Siman R, Cui H, Wewerka SS, Hamel L, Smith DH, Zwank MD. Serum SNTF, a Surrogate Marker of Axonal Injury, Is Prognostic for Lasting Brain Dysfunction in Mild TBI Treated in the Emergency Department. Front Neurol. 2020 Apr;11:249.

120. Gan ZS, Stein SC, Swanson R, Guan S, Garcia L, Mehta D, et al. Blood Biomarkers for Traumatic Brain Injury: A Quantitative Assessment of Diagnostic and Prognostic Accuracy. Front Neurol. 2019 Apr;10:446.

121. Siman R, Giovannone N, Hanten G, Wilde EA, McCauley SR, Hunter JV, et al. Evidence That the Blood Biomarker SNTF Predicts Brain Imaging Changes and Persistent Cognitive Dysfunction in Mild TBI Patients. Front Neurol. 2013 Nov;4:190.

122. Smith DH, Hicks R, Povlishock JT. Therapy Development for Diffuse Axonal Injury. J Neurotrauma. 2013 Mar;30(5):307–23.

123. Amoo M, O’Halloran PJ, Henry J, Ben Husien M, Brennan P, Campbell M, et al. Permeability of the Blood–Brain Barrier after Traumatic Brain Injury: Radiological Considerations. J Neurotrauma. 2022 Jan;39(1–2):20–34.

124. Bellaver B, Povala G, Ferreira PCL, Ferrari-Souza JP, Leffa DT, Lussier FZ, et al. Astrocyte reactivity influences amyloid-β effects on tau pathology in preclinical Alzheimer’s disease. Nat Med. 2023 Jul;29(7):1775–81.

125. Yang Z, Arja RD, Zhu T, Sarkis GA, Patterson RL, Romo P, et al. Characterization of Calpain and Caspase-6-Generated Glial Fibrillary Acidic Protein Breakdown Products Following Traumatic Brain Injury and Astroglial Cell Injury. Int J Mol Sci. 2022 Aug;23(16):8960.

126. Okonkwo DO, Yue JK, Puccio AM, Panczykowski DM, Inoue T, McMahon PJ, et al. GFAP-BDP as an acute diagnostic marker in traumatic brain injury: results from the prospective transforming research and clinical knowledge in traumatic brain injury study. J Neurotrauma. 2013 Sep;30(17):1490–7.

127. Kochanek PM, Bramlett H, Dietrich WD, Dixon CE, Hayes RL, Povlishock J, et al. A novel multicenter preclinical drug screening and biomarker consortium for experimental traumatic brain injury: operation brain trauma therapy. J Trauma. 2011 Jul;71(1 Suppl):S15–24.

128. Lafrenaye A, Mondello S, Povlishock J, Gorse K, Walker S, Hayes R, et al. Operation Brain Trauma Therapy: An Exploratory Study of Levetiracetam Treatment Following Mild Traumatic Brain Injury in the Micro Pig. Front Neurol. 2021 Jan;11:586958.

129. Zanier ER, Bertani I, Sammali E, Pischiutta F, Chiaravalloti MA, Vegliante G, et al. Induction of a transmissible tau pathology by traumatic brain injury. Brain. 2018 Sep;141(9):2685–99.

130. Kondo A, Shahpasand K, Mannix R, Qiu J, Moncaster J, Chen CH, et al. Antibody against early driver of neurodegeneration cis P-tau blocks brain injury and tauopathy. Nature. 2015 Jul;523(7561):431–6.

131. Zoltewicz JS, Mondello S, Yang B, Newsom KJ, Kobeissy F, Yao C, et al. Biomarkers Track Damage after Graded Injury Severity in a Rat Model of Penetrating Brain Injury. J Neurotrauma. 2013 Jul;30(13):1161–9.

132. Papa L, Zonfrillo MR, Welch RD, Lewis LM, Braga CF, Tan CN, et al. Evaluating glial and neuronal blood biomarkers GFAP and UCH-L1 as gradients of brain injury in concussive, subconcussive and non-concussive trauma: a prospective cohort study. BMJ Paediatr Open. 2019 Aug;3(1):e000473.

133. Levin HS, Temkin NR, Barber J, Nelson LD, Robertson C, Brennan J, et al. Association of Sex and Age With Mild Traumatic Brain Injury-Related Symptoms: A TRACK-TBI Study. JAMA Netw Open. 2021 Apr;4(4):e213046.

134. Bazarian JJ, Biberthaler P, Welch RD, Lewis LM, Barzo P, Bogner-Flatz V, et al. Serum GFAP and UCH-L1 for prediction of absence of intracranial injuries on head CT (ALERT-TBI): a multicentre observational study. Lancet Neurol. 2018 Sep;17(9):782–9.

135. Steyerberg EW, Wiegers E, Sewalt C, Buki A, Citerio G, De Keyser V, et al. Case-mix, care pathways, and outcomes in patients with traumatic brain injury in CENTER-TBI: a European prospective, multicentre, longitudinal, cohort study. Lancet Neurol. 2019 Oct;18(10):923–34.

136. Magatti M, Pischiutta F, Ortolano F, Pasotti A, Caruso E, Cargnoni A, et al. Systemic immune response in young and elderly patients after traumatic brain injury. Immun Ageing. 2023 Aug;20(1):41.

137. Moro F, Pischiutta F, Portet A, Needham EJ, Norton EJ, Ferdinand JR, et al. Ageing is associated with maladaptive immune response and worse outcome after traumatic brain injury. Brain Commun. 2022 Mar;4(2):fcac036.

