## Supplementary table 1 for "Translating from mice to humans: using preclinical blood-based biomarkers for the prognosis and treatment of traumatic brain injury"

Supplementary Table 1: Quality Score of studies

| Reference | peer rev | temperature control during surgery / treatment | surgery randomization | blinded outcome assasment | strain, age and sex reporting | sample size calulation | welfare regulation | conflict of interest | SUM |
|---|---|---|---|---|---|---|---|---|---|
| Ahmed et al., 2013 | ✓ | | ✓ | | ✓ | | ✓ | ✓ | 5 |
| Ahmed et al., 2015 | ✓ | | | | ✓ | | ✓ | ✓ | 4 |
| Aleem et al., 2022 | ✓ | ✓ | | | ✓ | | ✓ | | 4 |
| Arun et al., 2013 | ✓ | | | | ✓ | | ✓ | ✓ | 4 |
| Bashir et al., 2020 | ✓ | | ✓ | ✓ | ✓ | | ✓ | ✓ | 6 |
| Boucher et al., 2022 | ✓ | ✓ | ✓ | ✓ | ✓ | | ✓ | ✓ | 7 |
| Boutte´ et al., 2016 | ✓ | ✓ | | | ✓ | | ✓ | ✓ | 5 |
| Bramlett et al., 2016 | ✓ | ✓ | ✓ | ✓ | ✓ | | ✓ | ✓ | 7 |
| Browning et al., 2016 | ✓ | ✓ | ✓ | ✓ | ✓ | | ✓ | ✓ | 7 |
| Button et al., 2021 | ✓ | | | ✓ | ✓ | | ✓ | ✓ | 5 |
| Candy et al., 2017 | ✓ | | ✓ | ✓ | ✓ | | ✓ | ✓ | 6 |
| Cheng et al., 2018 | ✓ | | | | ✓ | | ✓ | ✓ | 4 |
| Cheng et al., 2023 | ✓ | | ✓ | | ✓ | | ✓ | ✓ | 5 |
| Çikriklar et al., 2013 | ✓ | | | | ✓ | | | ✓ | 3 |
| Çikriklar et al., 2016 | ✓ | | | | ✓ | | ✓ | | 3 |
| Dickstein et al.., 2021 | ✓ | | ✓ | ✓ | ✓ | | ✓ | ✓ | 6 |
| Diomede et al., 2023 | ✓ | | ✓ | ✓ | ✓ | ✓ | ✓ | ✓ | 7 |
| Dixon et al., 2016 | ✓ | ✓ | ✓ | ✓ | ✓ | | ✓ | ✓ | 7 |
| Ekici et al., 2014 | ✓ | | | | ✓ | | ✓ | ✓ | 4 |
| Gao et al., 2020 | ✓ | ✓ | | | ✓ | | ✓ | ✓ | 5 |
| Graham et al., 2021 | ✓ | ✓ | ✓ | ✓ | ✓ | | ✓ | ✓ | 7 |
| Heiskanen et al., 2022 | ✓ | | | | ✓ | | ✓ | ✓ | 4 |
| Hiskens et al., 2021 | ✓ | ✓ | ✓ | ✓ | ✓ | ✓ | ✓ | ✓ | 8 |
| Huang et al., 2015 | ✓ | ✓ | ✓ | | ✓ | | ✓ | | 5 |
| Jha et al., 2020 | ✓ | ✓ | ✓ | ✓ | ✓ | | ✓ | ✓ | 7 |
| Kamnaksh et al., 2012 | ✓ | | | | ✓ | | ✓ | ✓ | 4 |
| Kim et al., 2022 | ✓ | | ✓ | | | | ✓ | ✓ | 4 |
| Kmetova et al., 2022 | ✓ | | ✓ | | ✓ | ✓ | ✓ | ✓ | 6 |
| Kobeissy et al., 2022 | ✓ | | | | ✓ | | ✓ | ✓ | 4 |
| Koza et al., 2023 | ✓ | | | ✓ | ✓ | | ✓ | ✓ | 5 |
| Li et al., 2015 | ✓ | | | | ✓ | | ✓ | ✓ | 4 |
| Li et al., 2020 | ✓ | | ✓ | ✓ | ✓ | | | ✓ | 5 |
| Li et al., 2021 | ✓ | | ✓ | | ✓ | | ✓ | ✓ | 5 |
| Liang et al., 2017 | ✓ | | | ✓ | ✓ | | ✓ | ✓ | 5 |
| Liliang et al., 2010 | ✓ | | | | ✓ | | ✓ | ✓ | 4 |
| Liu et al., 2014 | ✓ | | ✓ | | ✓ | | ✓ | | 4 |
| Liu et al., 2010 | ✓ | ✓ | | | ✓ | | ✓ | ✓ | 5 |
| Mondello et al., 2016 | ✓ | ✓ | ✓ | ✓ | ✓ | | ✓ | ✓ | 7 |
| Moro et al., 2022 | ✓ | ✓ | ✓ | ✓ | ✓ | | ✓ | ✓ | 7 |
| Morris et al., 2019 | ✓ | | | | ✓ | | ✓ | ✓ | 4 |
| Mountney et al., 2016 | ✓ | ✓ | ✓ | ✓ | ✓ | | ✓ | ✓ | 7 |
| O'Brien et al., 2021 | ✓ | | ✓ | ✓ | ✓ | | ✓ | ✓ | 6 |
| O'Brien et al., 2023 | ✓ | | ✓ | ✓ | ✓ | | ✓ | ✓ | 6 |
| Osier et al., 2021 | ✓ | ✓ | ✓ | ✓ | ✓ | | ✓ | ✓ | 7 |
| Pham et al., 2021 | ✓ | | ✓ | | ✓ | | ✓ | ✓ | 5 |

| Reference | peer rev | temperature control during surgery / treatment | surgery randomization | blinded outcome assassment | strain, age and sex reporting | sample size calulation | welfare regulation | conflict of interest | SUM |
|---|---|---|---|---|---|---|---|---|---|
| Pierce et al., 2017 | ✓ | ✓ | ✓ | ✓ | ✓ | | ✓ | ✓ | 7 |
| Plog et al., 2015 | ✓ | | ✓ | | ✓ | | ✓ | ✓ | 5 |
| Pozdnyakov et al., 2019 | ✓ | | | | ✓ | | ✓ | ✓ | 4 |
| Robinson et al., 2016 | ✓ | | ✓ | ✓ | ✓ | ✓ | ✓ | ✓ | 7 |
| Rostami et al., 2012 | ✓ | | | | ✓ | | ✓ | ✓ | 4 |
| Rubenstein et al., 2019 | ✓ | ✓ | ✓ | ✓ | ✓ | | ✓ | ✓ | 7 |
| Rubenstein et al., 2015 | ✓ | | | | ✓ | | ✓ | ✓ | 4 |
| Rubenstein et al., 2017 | ✓ | | | | ✓ | | ✓ | ✓ | 4 |
| Saletti et al., 2023 | ✓ | ✓ | ✓ | ✓ | ✓ | ✓ | ✓ | ✓ | 8 |
| Scrimgeour et al., 2021 | ✓ | ✓ | ✓ | | ✓ | ✓ | ✓ | ✓ | 7 |
| Shear et al., 2016 | ✓ | ✓ | ✓ | ✓ | ✓ | | ✓ | ✓ | 7 |
| Siahaan et al., 2018 | ✓ | | ✓ | | ✓ | | ✓ | ✓ | 5 |
| Stemper et al., 2022 | ✓ | | | | ✓ | | ✓ | ✓ | 4 |
| Svetlov et al., 2012 | ✓ | | | | | | ✓ | ✓ | 3 |
| Svetlov et al., 2010 | ✓ | | | | | | ✓ | ✓ | 3 |
| Tadepalli et al., 2020 | ✓ | ✓ | | | ✓ | | ✓ | ✓ | 5 |
| Thau-Zuchman et al., 2020 | ✓ | ✓ | | ✓ | ✓ | ✓ | ✓ | ✓ | 7 |
| Tomita et al., 2020 | ✓ | ✓ | ✓ | | ✓ | ✓ | ✓ | ✓ | 7 |
| VandeVord et al., 2016 | ✓ | | ✓ | | ✓ | | ✓ | ✓ | 5 |
| Wallen et al., 2022 | ✓ | | ✓ | ✓ | ✓ | ✓ | ✓ | ✓ | 7 |
| Wei et al., 2015 | ✓ | | ✓ | ✓ | ✓ | | ✓ | ✓ | 6 |
| Wong et al., 2021 | ✓ | | ✓ | ✓ | ✓ | | ✓ | ✓ | 6 |
| Yang et al., 2013b | ✓ | | | | ✓ | | ✓ | ✓ | 4 |
| Yang et al., 2015 | ✓ | | ✓ | | ✓ | | ✓ | ✓ | 5 |
| Yao et al 2008 | ✓ | | | | ✓ | | ✓ | ✓ | 4 |
| Zoltewicz et al., 2013 | ✓ | ✓ | | | ✓ | | ✓ | ✓ | 5 |
